# Combined generalist and host-specific transcriptional strategies enable host generalism in the fungal pathogen *Botrytis cinerea*

**DOI:** 10.1101/2025.07.24.666639

**Authors:** Ritu Singh, Anna Jo Muhich, Cloe Tom, Jack McMillan, Karishma Srinivas, Lucca Faieta, Celine Caseys, Daniel J Kliebenstein

## Abstract

How generalist pathogens infect phylogenetically diverse hosts remains a central question in plant-pathogen biology. In particular, the extent to which broad host range is enabled by genetic variation versus transcriptional plasticity is unclear. To investigate how variation and plasticity contribute to generalism, we studied the generalist necrotrophic fungus *Botrytis cinerea* that infects more than 1,500 plant species. Using a cross-infection matrix of 72 *B. cinerea* isolates infected on 57 plant genotypes distributed across 15 eudicot species, we identified general and host-dependent fungal components of lesion formation. Transcriptome profiling at 48 hours post-inoculation revealed two distinct pathogen gene modules: (1) a set of general lesion-associated genes enriched in primary metabolism, showing similar expression across hosts but varied among isolates; and (2) a set of high-entropy, host specific-inducible genes, organized into distinct co-regulated modules that respond dynamically to specific host cues. Both gene sets were genomically dispersed, lacking structural clustering, and were under different levels of selective constraints. Our results demonstrate that *B. cinerea* employs a modular transcriptional strategy that integrates a core metabolic program along with a plastic, host-responsive regulatory network to achieve broad host colonization. This study presents the most comprehensive cross-species co-transcriptomic dataset to date for any fungal phytopathogen, highlighting transcriptional plasticity as a key mechanism underlying generalism in plant–fungal interactions. Moreover, the identification of conserved fungal gene targets across diverse hosts offers a foundation for developing broad-spectrum resistance strategies in multiple crops.

## Introduction

How and why some pathogens can infect hundreds of hosts while others are constrained to a single host remains an unfilled gap in our understanding of host-pathogen interactions and adaptability. A pathogen’s host range dynamic can be shaped by a complex interplay of genetic factors, environmental heterogeneity, and ecological interactions, which collectively drive host shifts, range expansions, or contractions. Furthermore, classical coevolutionary models, especially the gene-for-gene model, explain specialization through tight associations between resistance genes and pathogen effectors/virulence factors (Anderson *et al*., 2010; Flor, 1971; Jones and Dangl, 2006; Pieterse *et al*., 2012; Sánchez-Vallet *et al*., 2018). While these models provide a mechanistic understanding of pathogen specialization, they offer less insight into how generalist pathogens overcome the diverse barriers spread across hosts to infect a diverse range of hosts.

Several genetic mechanisms have been suggested to enable pathogens’ host range expansion, including mutation, hybridization, genome rearrangement, and horizontal gene transfer (Parrish *et al*., 2008; Pepin *et al*., 2010; Stukenbrock and Bataillon, 2012). While these gradual processes are important, they don’t fully explain the ability of some pathogens to rapidly shift hosts. Recent work has begun to link these broad host shifts to phenotypic plasticity, the ability of a single genotype to produce different phenotypes in response to environmental or biotic cues (Hu and Barrett, 2017). Three conceptual models describe how plasticity may facilitate host shifts: (1) ecological fitting, where pathogens exploit conserved traits across related hosts; (2) adaptive plasticity, in which induced traits enhance fitness on novel hosts; and (3) nonadaptive plasticity, where initial maladaptive responses expose cryptic genetic variation that may promote long-term adaptation (De Fine Licht, 2018). While these various plasticity models have been studied in modulating abiotic stress responses (Alster *et al*., 2021; Fox *et al*., 2019; Kronholm *et al*., 2016), their relevance to host-pathogen interactions, where both partners dynamically influence each other’s biology, remains elusive.

Unlike viruses and bacteria, which readily adapt to hosts via high mutation rates, eukaryotic pathogens may rely more on transcriptional reprogramming to facilitate broad host colonization (Beekman and Ene, 2020). For instance, in the ectoparasite crustacean *Tracheliastes polycolpus*, transcriptional plasticity enabled transcriptomic shifts across native and novel fish hosts (Mathieu-Bégné *et al*., 2022). In the plant pathogen *Fusarium oxysporum*, infection on nonvascular liverworts involved upregulation of conserved effector genes, while lineage-specific effectors associated with angiosperm infection remained silent (Srivastava *et al*., 2025). Similarly, *Fusarium virguliforme* exhibits host-dependent transcriptional programs that differ between the pathogenic lifestyle on soybean and endophytic colonization on maize (Baetsen-Young *et al*., 2021). Co-transcriptomic analyses of a single isolate of the generalist fungal pathogen *Sclerotinia sclerotiorum* across six host species revealed both generalist and polyspecialist transcriptional strategies, with distinct gene expression programs deployed depending on the host, suggesting that plasticity can enable a flexible generalist infection strategy (Kusch *et al*., 2022).

While these studies highlight the potential role of transcriptional plasticity in facilitating infection on different hosts, most have focused on either single pathogen isolates or a limited set of hosts. Thus, it remains unclear how genetic diversity in the pathogen populations modulates transcriptional plasticity across phylogenetically diverse hosts. Generalist pathogens may employ overlapping strategies, including standing genetic variation, core and lineage-specific virulence programs, and transcriptional plasticity, to overcome diverse host defenses. Dissecting the relative contribution of each mechanism requires a broad, systematic, and phylogenetically informed approach that captures both host diversity and pathogen intraspecific variation.

*Botrytis cinerea* provides a powerful model to address these questions. This broad host-range necrotrophic fungus infects over 1,500 plant species across angiosperms, gymnosperms, and mosses, with a notable preponderance among eudicots (Elad and Fillinger, 2016; Singh *et al*., 2024). While *B. cinerea* exhibits extensive standing genetic diversity and polygenic virulence architecture (Bi *et al*., 2023; Caseys and Kliebenstein, 2025; Krishnan *et al*., 2023; Mbengue *et al*., 2016; Williamson *et al*., 2007), the extent to which *B. cinerea* transcriptionally adapts to diverse hosts remains unresolved. A key advantage enabling this question is that *B. cinerea* can be readily infects many host species, and its transcripts can be simultaneously measured from host– pathogen RNA-seq data, enabling efficient and scalable co-transcriptome profiling (Zhang *et al*., 2019).

In this study, we use the *B. cinerea*-eudicot pathosystem to investigate how phenotypic and transcriptional plasticity, alongside genetic diversity of both host and pathogen, contribute to the generalist lifestyle of *B. cinerea*. We assayed lesion development across 15 eudicot host species spanning eight orders, using four genotypes per species and 72 genetically diverse *B. cinerea* isolates, capturing a wide range of natural variation (Atwell *et al*., 2018; Caseys *et al*., 2021; Corwin *et al*., 2016; Soltis *et al*., 2020; Zhang *et al*., 2017; Zhang *et al*., 2019). To investigate transcriptional dynamics during infection, we built a co-transcriptome dataset with 10 representative host species from six eudicot orders using the same panel of 72 isolates. This large-scale, phylogenetically informed design enables us to explore how host evolutionary distance and pathogen genetic diversity shape infection outcomes and fungal gene expression programs. By integrating phenotypic and transcriptomic data across diverse host–pathogen combinations, we aim to uncover how plasticity and genetic diversity drive broad host colonization.

## Results

### Comparative Lesion Development

To investigate the interaction of a *B. cinerea* population across diverse host species, we infected 57 plant genotypes representing 15 eudicots across eight taxonomic orders with 72 *B. cinerea* isolates. This approach generated an infectivity matrix of 26,718 independent lesions at 72 hours post-inoculation (hpi) **(Figure 1; Table S1)**. Notably, lesions typically became visible only after 48 hpi in most species, preventing digital image analysis of earlier time points. To understand how the lesions further progressed from 72 to 96 hpi, we calculated the relative growth rate (RGR = [lesion 96 hpi – lesion 72 hpi)/ 24)] of each isolate. The RGR showed a linear relationship with lesion size at 72 hpi across all species, suggesting that lesion size at 72 hpi is a reliable proxy for *B. cinerea* growth potential. Notably, the slope of the relationship varies among species **(Figure S1A)**. Parsley, Pepper, Cowpea, Bean, and Cucumber exhibited faster growth from 72 to 96 hpi, whereas Chard and Celery showed slower progression, likely reflecting differences in post-penetration host responses (**Figure S1**). The rank order of the isolate virulence at 72 hpi versus 96 hpi did not change within a host, as expected for a linear relationship (**Figure S1B**). Therefore, the lesion area at 72 hpi was selected as the optimal time point for investigating host-pathogen interaction dynamics while avoiding any potential constraints on lesion development caused by tissue availability.

**Figure 1:**
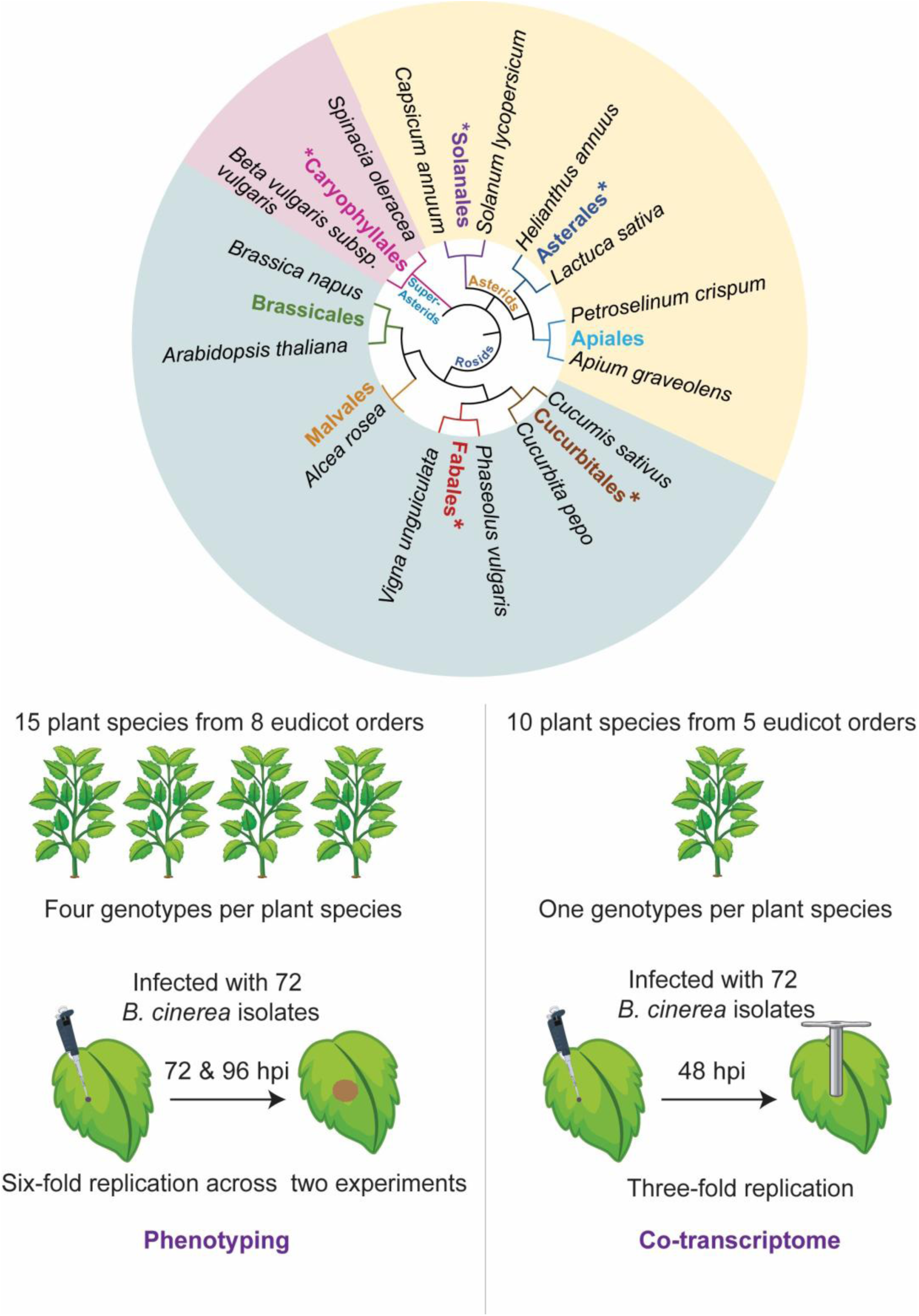
Experimental design for lesion phenotyping and co-transcriptome analysis. We tested the virulence of 72 *B. cinerea* isolates across 15 eudicot plant species from 8 eudicot orders, selected to represent a balanced phylogenetic distribution. Each plant species included four different genotypes used for phenotyping assays. For lesion phenotyping, leaves were inoculated and lesion sizes measured at 72- and 96-hours post-inoculation (hpi) with six-fold replication across two independent experiments. A subset of 10 plant species from 5 eudicot orders (marked with **\*** in the phylogenetic tree) were selected for co-transcriptomic profiling. These species were inoculated with the same 72 isolates, and leaf tissue was harvested at 48 hpi for co-transcriptome analysis, with three-fold replication. Leaf disks were collected using a cork borer of the same size to ensure consistency in tissue quantity across samples.

### Quantitative variation in host susceptibility to *B. cinerea* across eudicot plant species

To assess the contribution of host and pathogen genetic diversity to variation in lesion size, we used a linear mixed model within individual plant species that accounted for experimental design variables. These experimental factors explained a significant proportion of variance in nearly all species (**Figure 2A**). Within each individual host species, genetic variation among the 72 *B. cinerea* isolates explained the largest proportion of lesion size variance, ranging from 15% to 45% of the total (**Figure 2A**). The interaction between host genotype and isolate contributed an additional 5% to 11% of the total variance, while host genotype alone explained a smaller but statistically significant fraction (0.6% to 6%). This indicates that within species, pathogen genetic diversity is a primary driver of lesion size variation, with host genotype playing a smaller but significant role.

**Figure 2.**
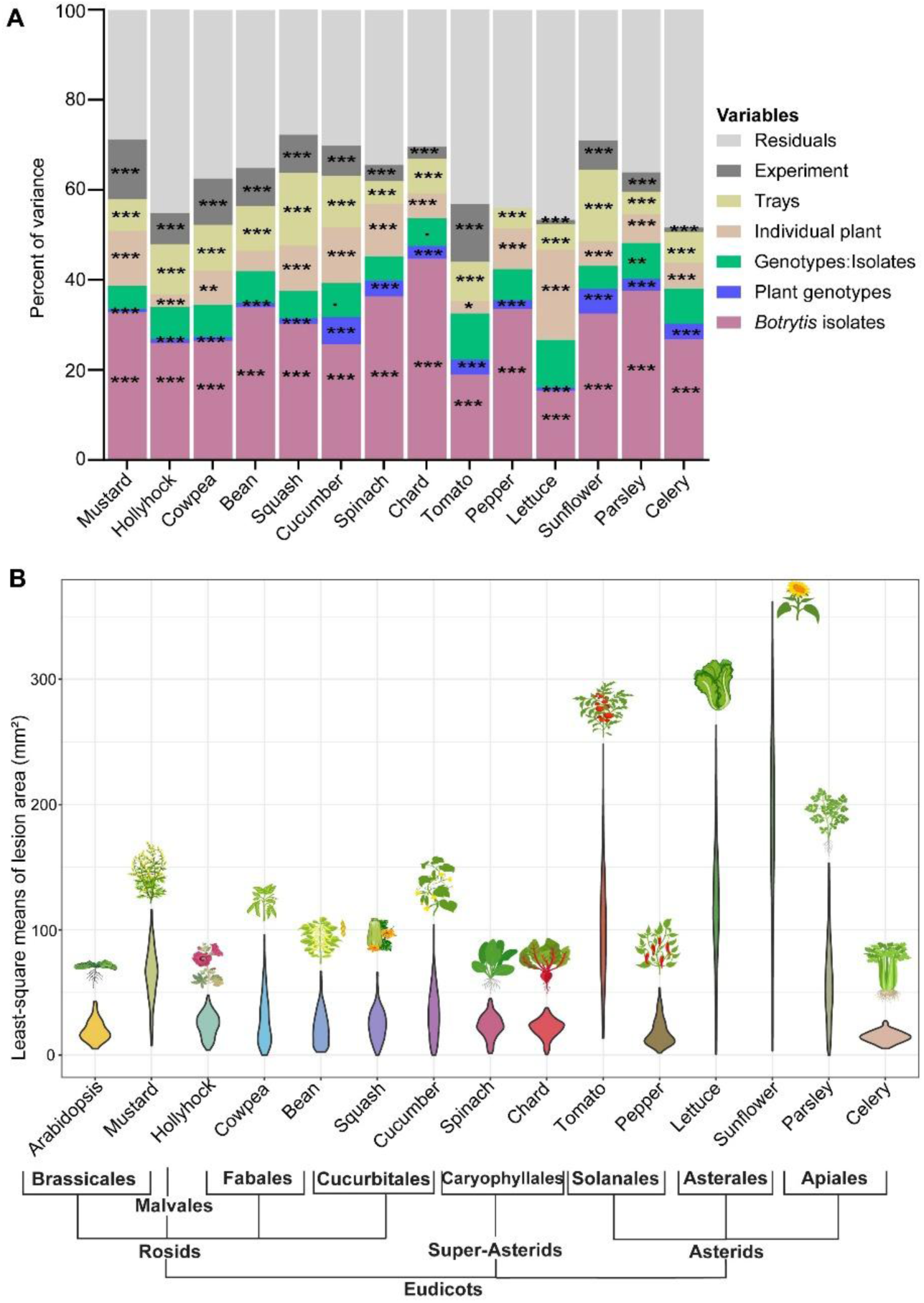
Quantitative variation in host susceptibility to *B. cinerea* across 15 eudicot species at 72 hours post inoculation (hpi). **(A)** The partitioning of experimental and genotypic variance within individual plant species using linear mixed models of lesion area at 72 hpi is shown. Stacked bar plots show the proportion of total variance in lesion size explained by various experimental factors. Fixed effects include *B. cinerea* isolate (purple), plant genotype (blue), and their interaction (green). Random effects include experimental design factors; individual plant, tray, and independent experiment (beige and gray tones). The model residual is shown in gray. Asterisks indicate the statistical significance of each variance component (\**p* < 0.05, \*\**p* < 0.01, \*\*\**p* < 0.001). **(B)** Violin plots represent the distribution of average lesion area (mm²) across 15 eudicot plants infected with 72 *B. cinerea* isolates. A non-scaled phylogenetic tree illustrates the evolutionary relationships among species. Illustrations of representative plant species are shown above each violin plot.

Comparing lesion area across the 15 host species showed substantial interspecific variation. Celery exhibited the smallest lesions, indicating high tolerance to *B. cinerea*, whereas Sunflower developed the largest, reflecting high susceptibility (**Figure 2B; Table S1)**. Genotypes within each plant species showed a similar range of lesion sizes (**Figure S2**). At a broader phylogenetic level, many species from the Rosids clade, such as *Arabidopsis*, Mustard, and Cowpea, displayed lower susceptibility than several Asterids, such as Lettuce and Sunflower. Species from the Caryophyllales (Spinach and Chard) were among the resistant species. These patterns highlight the diversity of the disease outcome and suggest a phylogenetic signal in *B. cinerea-*eudicot interactions across eudicots.

### Host-pathogen interaction patterns are predominantly shaped at the species level

To evaluate whether host-specific differences in lesion size reflect the hosts evolutionary relationships, we assessed how the eudicot phylogeny associates with *B. cinerea* lesion formation. A multi-host nested linear mixed model across the host phylogeny showed that the largest contribution to the percentage of variance in lesion area is explained by the orders (35%), followed by clades (23%), species (16%), and then genotypes (3%) (**Figure 3A**). When considering interaction with the pathogen, isolate-by-species and isolate-by-order both explain 5% of the total variance, with a minor contribution from isolate-by-clade (2%). *B. cinerea* isolates alone contributed 5% of the total variance. These findings suggest that while *B. cinerea* isolate variation is a major driver of lesion variation within species, across the host phylogeny; the diversity of disease outcome is primarily shaped by hosts and host-pathogen interactions.

**Figure 3.**
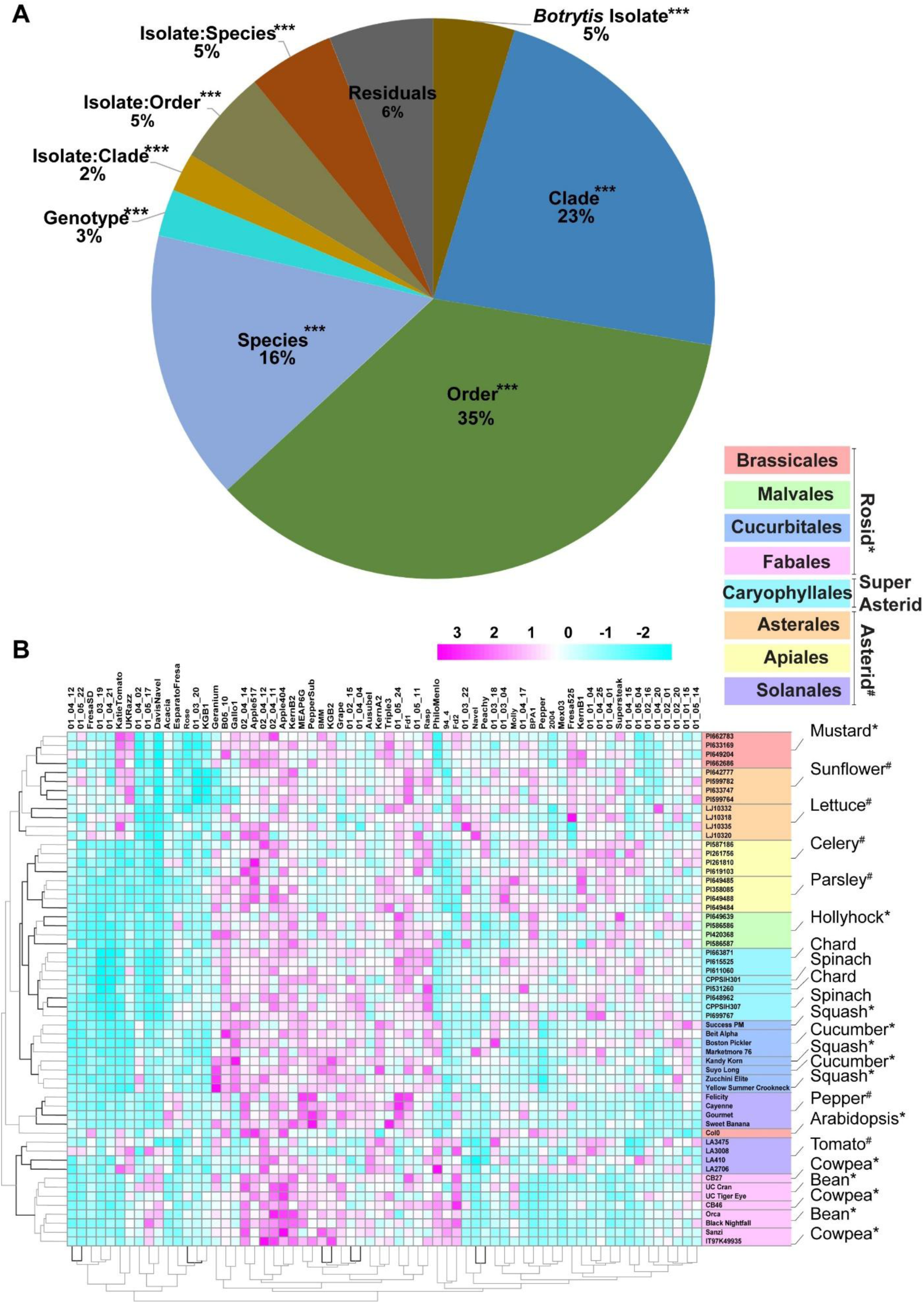
Plant susceptibility to *B. cinerea* shows species and order-specific patterns across the eudicot phylogeny. **(A)** Variance partitioning using a multi-host linear model to estimate the percentage of variance in lesion area explained by host phylogenetic levels (clade, order, species, and genotype), *B. cinerea* isolate, and their interactions. Host levels were nested according to taxonomic hierarchy. This model quantifies both host and pathogen contributions to observed lesion variation across the full dataset. Asterisks (\*\*\**p* < 0.001) indicate the statistical significance of each variance component. **(B)** Heatmap showing standardized (z-scored) least squares mean lesion areas from infection assays between 72 *B. cinerea* isolates (columns) and 61 plant genotypes (rows). Each row corresponds to a unique plant genotype, with genotype names followed by their respective species. All plant species are represented by four genotypes, except Arabidopsis, which has a single genotype (Col-0). Colored sidebars indicate plant taxonomic orders; species labeled with an **\*** belong to the Rosid clade, a **#** denotes Asterid, and unlabeled species belong to the Super-Asterid group. Dendrograms indicate hierarchical clustering of isolates and host genotypes, with bold branches denoting clusters with ≥ 95% bootstrap support.

To further explore this phylogenetic structure, we performed hierarchical clustering of lesion area across all host genotypes and *B. cinerea* isolates. Most of host genotypes clustered within their species, with notable exceptions in Bean, Cowpea, and one Chard genotype, where genotypes did not cluster together within the species, but within the order. (**Figure 3B**). Order-level grouping was also evident for most species, supporting a potential phylogenetic signal in lesion formation. However, the species and order level clustering collapsed at the clade level with Rosids and Asterids being interspersed (**Figure 3B**). While deeper taxonomic structures were statistically significant, their impact on disease outcome is limited compared to lineage-specific variation. These patterns suggest that *B. cinerea*-eudicot interactions are largely influenced by variation within or between closely related host species or rather than by deeper evolutionary branches.

### General and host-dependent lesion-forming potential in *B. cinerea*

The modest but statistically significant phylogenetic signal across all hosts suggests a host-independent virulence component across the *B. cinerea* isolates. To estimate such effects, we calculated the average lesion size of each isolate across all hosts. This metric, hereafter called general lesion, captures the lesion-forming potential of an isolate across all hosts. Additionally, we generated a variable called host-dependent lesion, which was calculated as the mean lesion area produced by each isolate within a single host species (see Materials and Methods) (Caseys and Kliebenstein, 2025; Corwin *et al*., 2016).

To assess the relationship between general and host-dependent lesion potential, we performed linear regression analyses. Across all host species, we observed significant positive correlations, indicating that isolates with higher general lesions tend to be more virulent across the individual hosts (**Figure 4A; Figure S3; Table S2**). The strength of this correlation varied widely among host species and orders. For example, in Caryophyllales (Spinach and Chard), general and host-dependent lesions were moderately correlated (*R²* = 0.54 and 0.46, respectively), indicating that disease outcomes in these hosts are driven by a combination of general lesion potential and some host-specific properties. In contrast, Fabales (Bean and Cowpea) showed weaker correlations (adjusted *R²* = 0.10 and 0.11, respectively), implying a greater contribution of host-specific interactions in shaping disease outcomes. These results indicate that while general aggressiveness partly contributes to lesion formation across all hosts, the disease outcome is heavily influenced by the host-specific aspects of the interaction.

**Figure 4:**
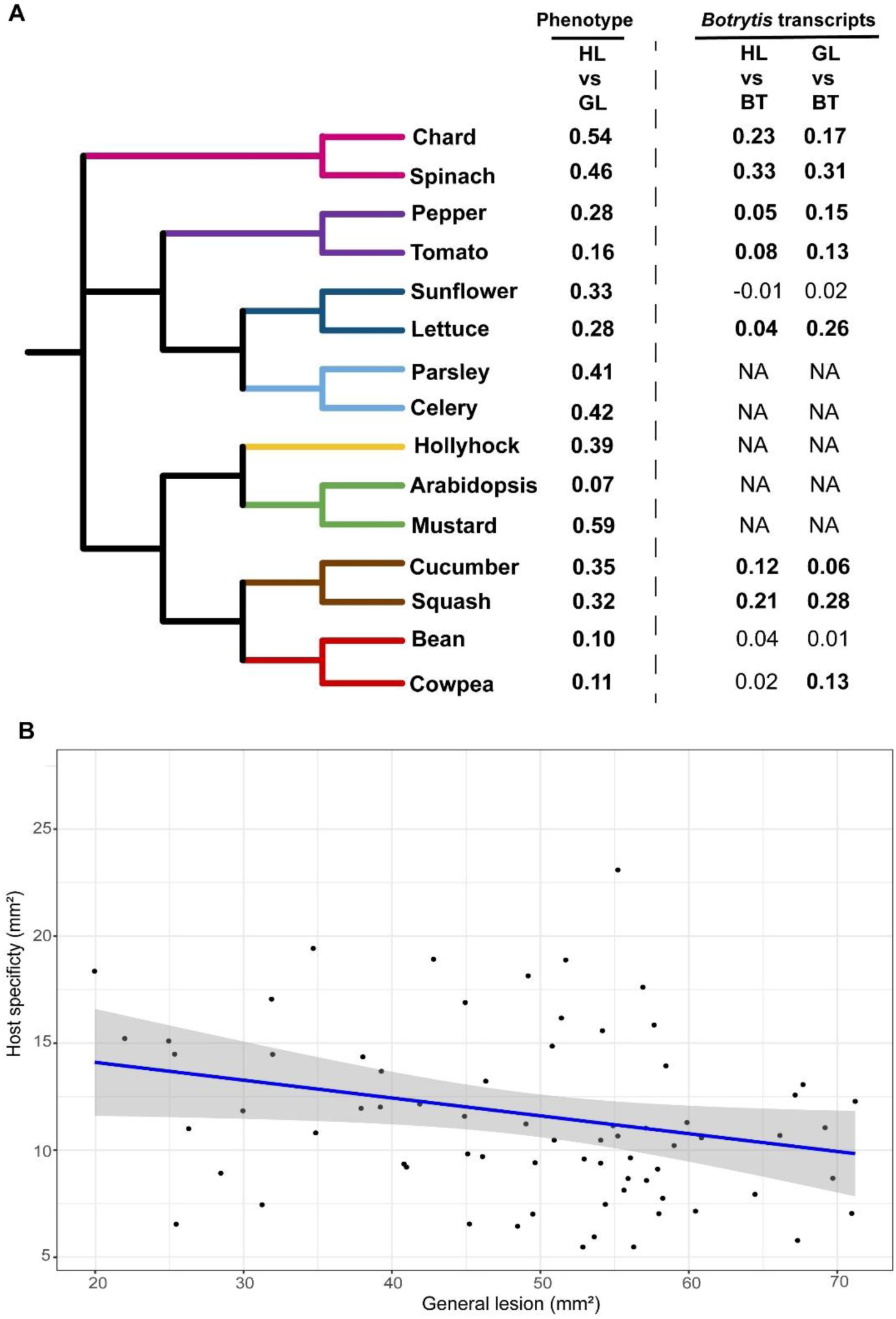
Phylogenetic patterns and descriptors of lesion size variation in *Botrytis cinerea* infections across eudicot hosts. **(A)** Phylogenetic tree of 15 eudicot plant species used in *B. cinerea* infection assays, color-coded by plant order. The adjacent table summarizes the coefficient of determination (*R²*) from linear regression models evaluating the association of different variables with lesion size outcomes. The GL (general lesion) variable represents the mean lesion size across all hosts for each isolate. This was recalculated independently for each species to exclude the focal species and avoid overestimation. HL (host-dependent lesion) for each isolate was calculated as the mean lesion on a given host, while correcting for the genotype-level variation. BT (*B. cinerea* transcript abundance) represents total *B. cinerea* transcript abundance at 48 hpi (a proxy for fungal biomass). “NA” indicates species for which transcriptomics were not performed. Bold values indicate statistically significant predictors (*p* < 0.05). Full model results and regression plots are shown in **Figures S3, S5, and S6 and Tables S2 and S4**. **(B)** Relationship across isolates between the host specificity and general lesion estimates (*R²* = 0.05, *p* = 0.04). Host specificity for each isolate was calculated as the total residual variance in lesion size across all hosts when comparing the isolates’ lesion size on each host against the general lesion estimate.

To estimate the host specificity potential of each isolate, we calculated the residuals from the linear regression between host-dependent and general lesions. These residuals capture isolate-by-host deviations not explained by the general aggressiveness of an isolate. A high host specificity score for an isolate indicates a larger deviation from the generalist predicted behavior across all the hosts, likely indicating host-specific lesion formation for that isolate. Comparing general lesion values to host specificity scores across the isolates shows a statistically significant but very small relationship between general lesion and host-specificity score (*R*² = 0.05, *p* = 0.04). This suggests that an isolate’s general aggressiveness across all host species is largely independent of its host-specific lesion formation capacity (**Figure 4B**).

To further test for host specialization based on the host from which the isolates were collected, we assessed whether the seven isolates collected from hosts tested in this assay [Brassica (*Ausubel*), Tomato (*Katie Tomato*, *KGB1*, *KGB2*, *Supersteak*, and *Triple3*), and Pepper (*Pepper*)] were more virulent on their respective source host. Across all hosts, none of these seven isolates prefer their respective source host; instead, they form bigger lesions on unrelated hosts (**Table S1**, **Figure S4**). These results suggest limited evidence for strict host specialization among our collection of isolates, aligning with previous studies (Ma and Michailides, 2005; Martinez *et al*., 2003; Rowe and Kliebenstein, 2007; Samuel *et al*., 2012; Soltis *et al*., 2020).

### Early *B. cinerea* Transcript Abundance Partially Predicts Lesion Development in a Host-Dependent Manner

The ability of nearly all the *B. cinerea* isolates to infect all the tested host plants suggests that each isolate likely can adjust to each host. As such, this indicates that *B. cinerea* may plastically modulate its transcriptome to enable lesion formation on the different hosts. To test this, we quantified how fungal gene expression changes across the host species, pathogen isolates, and their interaction. We conducted a co-transcriptome analysis at 48 hpi across ten eudicot species infected with 72 *B. cinerea* isolates (**Figure 1; Table S1 and S3**). As an initial analysis, we tested if early-stage fungal transcript abundance could serve as a predictive marker for subsequent lesion development. For this, we modeled the relationship between the total amount of *B. cinerea* transcripts at 48 hpi with the resulting lesion formed at 72 hpi across eudicot hosts using linear regression. We postulated that the proportion of fungal-mapped reads relative to total mapped reads may be a proxy for fungal biomass (Blanco-Ulate *et al*., 2014; Zhang *et al*., 2019). For most species, early fungal transcript abundance showed a weak but significant correlation with both host-dependent lesion and general lesion ability, with maximum *R²* values around 0.3 (**Figure 4A**, **Figures S5 and S6, Table S4**). These results suggest that early fungal gene expression partially connects with subsequent lesion formation as measured by lesion size.

### Genetic Variation in the Pathogen and Between Hosts Alter *B. cinerea* Transcriptome

To disentangle the contributions of host, pathogen, and their interaction to *B. cinerea* transcripts expression, we fit linear models for each gene across all host–pathogen combinations within a single model and estimated broad-sense heritability (*H²*) for each genetically variable component (Krishnan *et al*., 2023; Soltis *et al*., 2020). Across the full dataset, 11,415 fungal genes were expressed, of which 6,824, 1,826, and 2,768 genes were significantly influenced by genetic variation in the host, pathogen, and host-pathogen interaction, respectively (**Table S5**). These indicate that across the full dataset, the fungal transcripts were primarily influenced by the host and host–pathogen interactions rather than the pathogen alone (**Figure 5A**; average *H²*: host = 0.14, pathogen = 0.07, interaction = 0.10).

**Figure 5:**
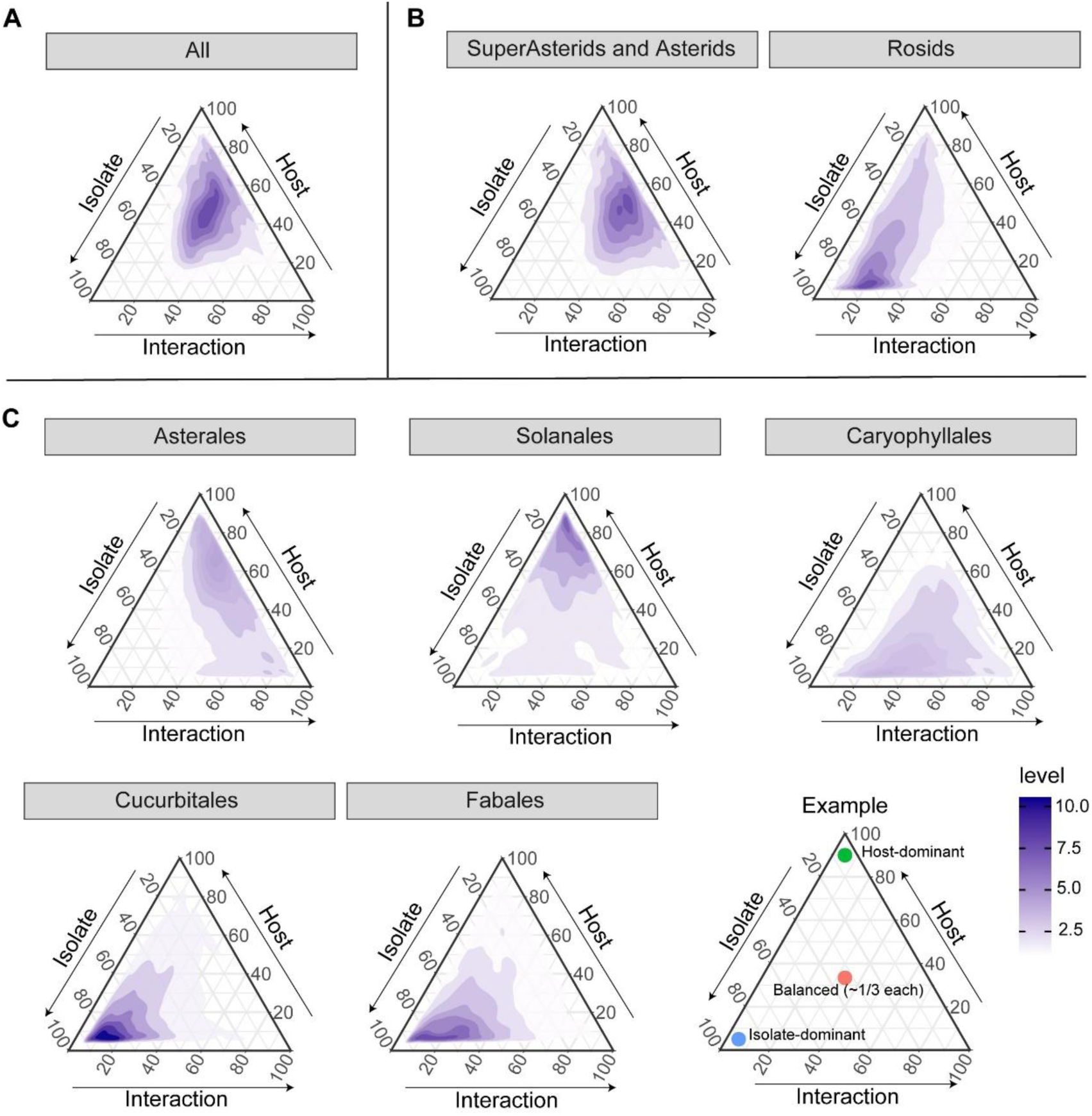
Differential genetic contribution of host and pathogen isolates to *B. cinerea* transcript variation across the full 10-host eudicot panel. Ternary plots illustrate the proportion of total heritability in *B. cinerea* gene expression explained by host plant species, *B. cinerea* isolate, and their interaction. Models were fitted separately at multiple phylogenetic levels: **(A)** across all host species (All), **(B)** within clades, and **(C)** within orders. The panel labeled Super-Asterids and Asterids integrates data from Asterales, Solanales, and Caryophyllales. Color contours (level) represent relative gene density. An annotated example ternary plot is provided to illustrate how to interpret component contributions.

To ascertain if these relative contributions vary across evolutionary lineages, we performed analyses at the clade and order levels (**Table S5; Figure 5B and C**). The patterns differed across groups: in Asterales, fungal gene expression was largely shaped by the host and interaction; in Solanales, all three components contributed evenly; and in Fabales and Cucurbitales, variation was primarily driven by the pathogen. Caryophyllales showed a unique pattern, with high heritability explained by both pathogen and host–pathogen interactions, but relatively less from the host alone. These together suggest that *B. cinerea* transcriptional responses genetically differ across isolates and are highly plastic to host species variation.

### Low-Entropy Genes Represent a Stable, Non-Virulence Core Transcriptome

The above analysis suggests that *B. cinerea* genes have diverse expression patterns across host plants. To better classify how the pathogen genes respond across hosts, we calculated the Shannon entropy for each gene using the average expression across all isolates for each host species (**Figure 6A**). Low-entropy genes are uniformly expressed across all hosts, while high-entropy genes exhibit host-specific expression patterns. Using the Jenks natural breaks algorithm, genes were classified into three entropy categories: low (n = 1), intermediate (n = 10,828), and high (n = 589) (**Table S6**).

**Figure 6.**
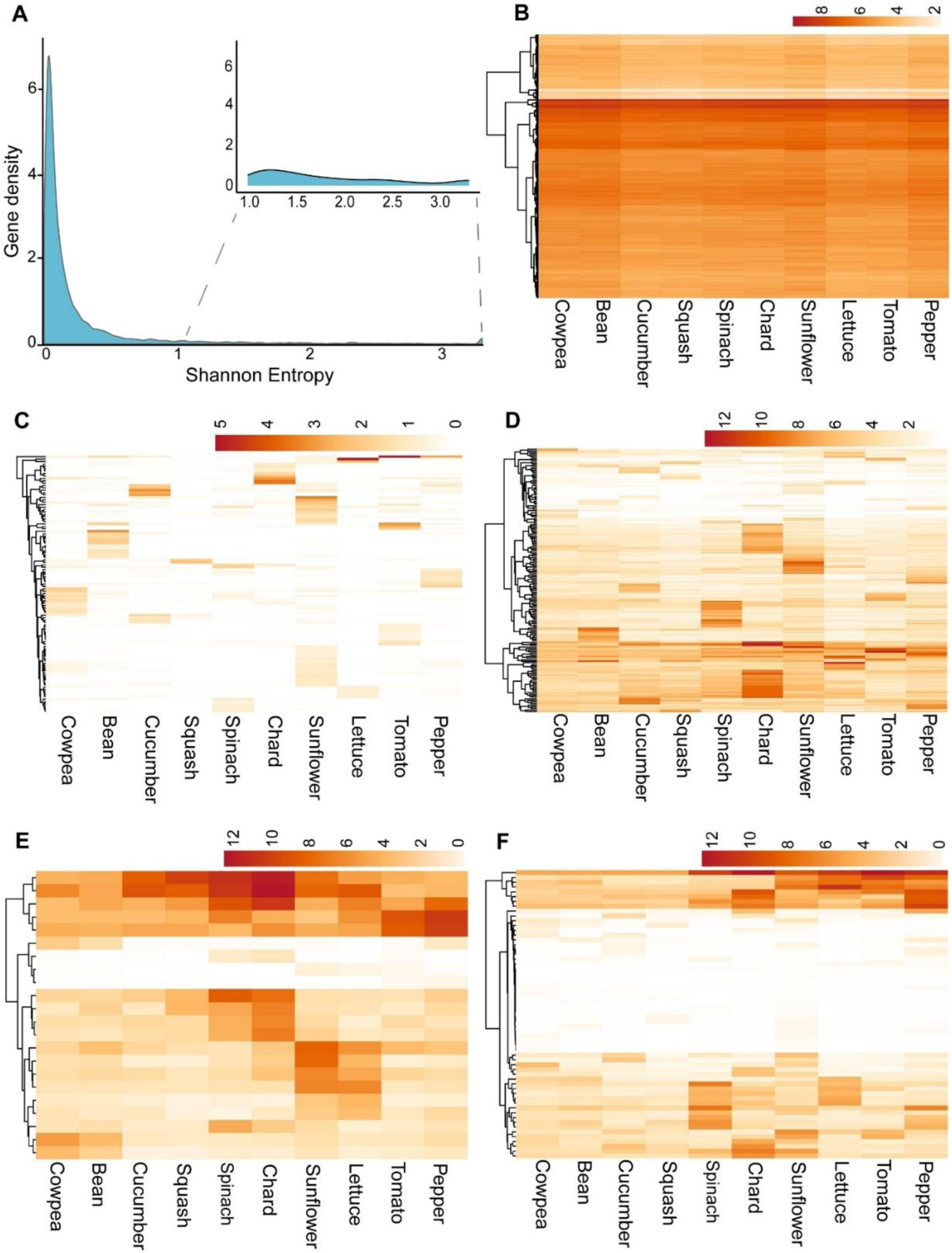
Conserved and host-specific *B. cinerea* gene expression across 10 eudicot hosts. **(A)** Density plot of Shannon entropy across *B. cinerea* genes calculated from host-averaged normalized CPM expression values. The inset zooms into the right tail of the distribution, highlighting high-entropy genes. **(B–F)** Heatmaps showing log₂ (CPM + 1) expression values for each of 10 host species, averaged across 72 *B. cinerea* isolates. Rows represent individual genes, hierarchically clustered based on expression patterns; columns represent plant host species, grouped by phylogenetic clade. The heatmap color scale represents an increasing expression intensity from low to high. A full isolate-level expression profile is provided in **Figures S7A, S14B, S14C, and S15**. **(B) Low entropy genes (lowest 500 genes):** Genes with consistent expression across most hosts, indicative of broadly conserved transcriptional activity. **(C and D) High-entropy genes with elevated expression in a single host (single-host-specific):** A gene was defined as specific to a single host if its expression in that host was ≥ 1 standard deviation (SD) above its mean expression across all hosts. **C.** 262 genes expressed in only a few hosts, with strong elevated expression (≥ 1 SD) in a single host. **D.** 172 genes expressed across all hosts, but with expression ≥ 1 SD higher in one host compared to others. **(E and F) High entropy genes (specific to two or three hosts):** Genes showing higher expression (≥ 1 SD above their mean expression across all hosts) in two or three hosts, compared to the remaining hosts, representing moderately specific expression. **E.** 22 genes with high expression in phylogenetically related hosts, suggesting order-specific regulatory responses. **F.** 60 genes are highly expressed in two or three phylogenetically unrelated hosts.

Since only one gene was assigned to a low-entropy class, we selected 500 genes with the lowest entropy values (≤ 0.015) for further analysis (**Table S7**). While these low-entropy genes are influenced by variation in host and isolate, their expression patterns are largely consistent across all samples (**Figure 6B, Figures S7A and S8**). Given their relatively stable expression pattern, we did not expect strong co-expression patterns among these genes. Consistent with this, co-expression network analysis revealed that only 38 out of 500 low-entropy genes (8%) grouped into a single module across the full isolate–host dataset (**Figure S7B**). Functional enrichment analysis of all low-entropy genes showed significant overrepresentation of kinase activity, DNA and protein binding, and core biosynthetic and regulatory processes (**Figure S7C-D**). Comparing these genes expression profiles *in planta* and *in vitro* on artificial media showed that all 500 genes were expressed under both conditions, with 90% showing a similar expression level between *in planta* and *in vitro* (**Figure S9A**). Correlating these genes expression with host-dependent lesion estimates identified no significant correlations (**Table S8**). As such, we hypothesize that these low-entropy genes represent a core expression program for *B. cinerea,* independent of the environment or plant.

### General Lesion-Associated Genes Are Enriched in Primary Metabolism

The majority (88%) of *B. cinerea* genes fell into the intermediate Shannon entropy category, indicative of broadly distributed but variable expression patterns across isolates and hosts (**Table S6)**. We hypothesized that this category may include candidates contributing to the variation in isolates’ general lesion-forming potential across hosts. To test this, we performed a linear model across all genes regardless of entropy and 72 *B. cinerea* isolates, correlating general lesion to fungal gene expression while controlling for host species and gene-by-host interaction effects. This approach identified 287 genes significantly associated with general lesion variation across all hosts and no host dependency (**Figure 7A; Table S9**). As expected, almost all of these genes belonged to the intermediate entropy category, with only two genes being identified from within the high-entropy genes. These general lesion-associated genes showed similar expression across all hosts but varied among isolates (**Figure S10**).

**Figure 7.**
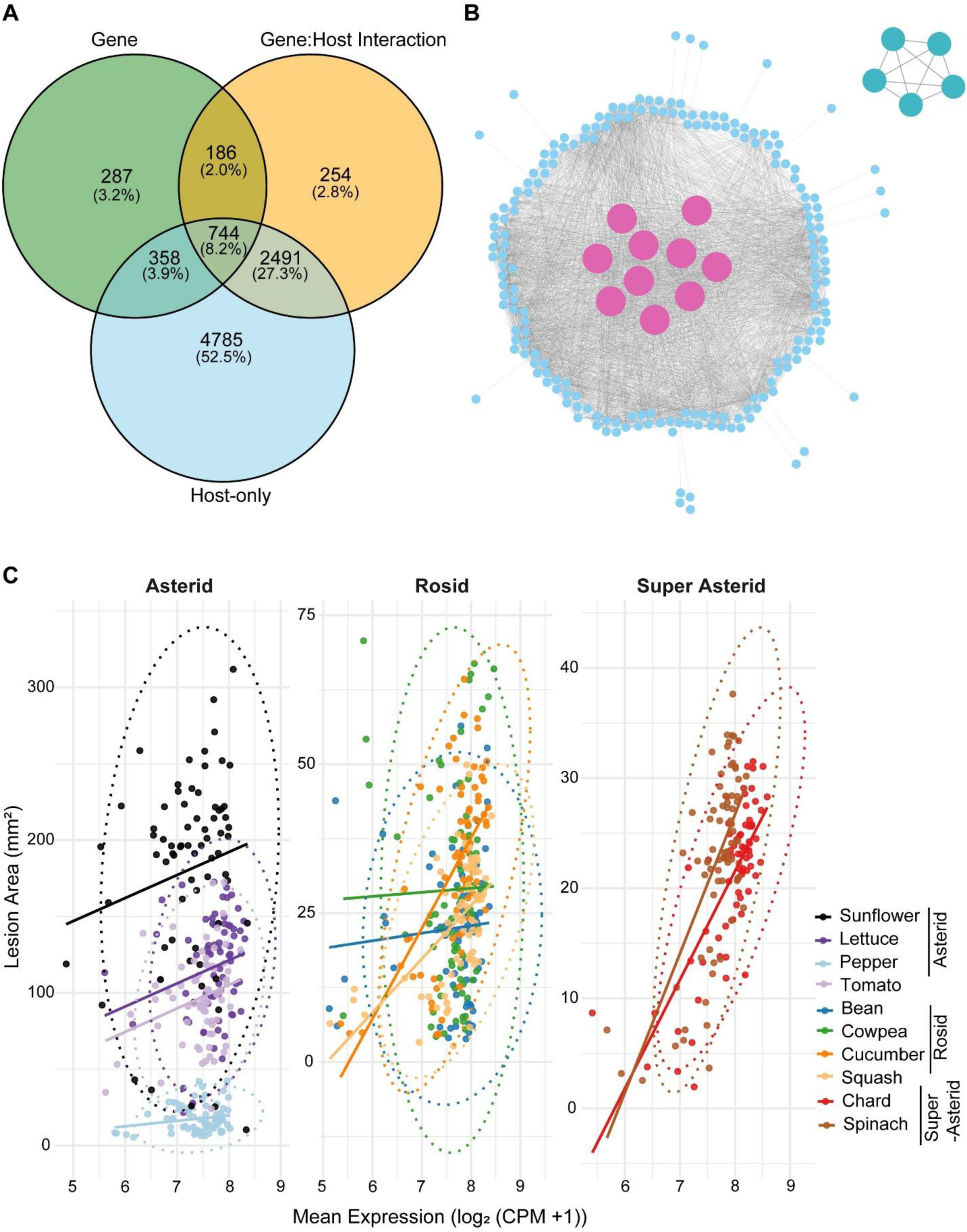
General lesion-associated module within *B. cinerea*. **(A)** Venn diagram showing the number of genes significantly associated with general lesion size (Gene), host identity (Host-only), and gene-by-host interaction (Gene:Host Interaction) based on individual gene linear models. A total of 287 genes were uniquely associated with the general lesion term and not confounded by host or interaction effects; these were retained as general lesion-associated candidates for further analysis. **(B)** Co-expression network of general lesion-associated genes constructed using absolute Pearson correlation (|R| ≥ 0.7) across all isolates and hosts. Nodes represent genes, and edges represent significant co-expression relationships. Pink nodes indicate hub genes within the main module. A second smaller module in the top right comprises five *Botcinic acid* (*BOA*; *Bcboa1*, *Bcboa2*, *Bcboa4*, *Bcboa6*, *Bcboa9*) genes. **(C)** Scatterplots showing the relationship between mean expression of general lesion-associated genes [log₂(CPM +1)] and host-dependent lesion area (mm²) across isolates for each host species. Hosts are grouped by phylogenetic clade (Asterid, Rosid, Super Asterid), and colored lines represent linear regressions for each species. Ellipses denote 95% confidence intervals for each host group. Each point represents an isolate. Rho and *p*-values for host-wise Spearman correlations are provided in **Table S8**.

To determine whether these genes were plant-induced or more broadly expressed, we compared their expression under *in vitro* and *in planta* conditions. All 287 genes were expressed in both conditions; however, 39% exhibited elevated expression *in planta*, 45% showed similar expression in both conditions, and 16% showed higher expression *in vitro* (**Figure S9B**). These results indicate that this gene module exhibits conditional plasticity, with many genes varying in expression depending on the growth environments. We next assessed the extent to which expression of these general lesion–associated genes were correlated with host-dependent lesion size. Significant positive correlations were observed in Chard (ρ = 0.69), Cucumber (ρ = 0.58), Spinach (ρ = 0.58), Squash (ρ = 0.48), Cowpea (ρ = 0.23), and Lettuce (ρ = 0.24), while the remaining four hosts showed no significant associations (**Table S8; Figure 7C**).

To identify potential mechanistic networks, we developed co-expression networks using these 287 genes expression across all the hosts. This identified two co-expression networks that differ by functional enrichment analysis. The large module, including 207 genes, was enriched for primary metabolic processes, and the smaller module contained five *Botcinic acid* (*BOA*) biosynthetic genes (*Bcboa1, Bcboa2, Bcboa4, Bcboa6, and Bcboa9*) (**Figure 7B; Figure S11**). This suggests that coordinated expression of core metabolic genes along with a selected phytotoxin pathway underlies variation in general lesion formation across isolates.

Interestingly, several canonical virulence factors, including cell wall–degrading enzymes (e.g., *PG1*), phytotoxins from the *Botrydial* (*BOT*) cluster, and other *BOA* cluster genes, were not identified among the general lesion-associated gene set. This is notable given previous hypotheses that such genes should be uniformly expressed across host species (Choquer *et al*., 2007; Mbengue *et al*., 2016; Newman and Derbyshire, 2020). Independent examination of these canonical genes expression profiles revealed that most of them were within the intermediate entropy category, but with moderate host-dependent variation in expression (**Figures S12-S13)**. This suggests that these genes expression are plastic in a host-dependent manner making their importance for virulence variable across all hosts.

### Host-specific transcriptional expression patterns

Next, to identify host-specific transcriptional responses, we analyzed 589 high-entropy genes that showed substantial expression variability across hosts. Using z-score normalization of log₂ (CPM + 1) values, we defined a gene as specific to a single host if its expression in that host was at least one standard deviation (SD) above its mean expression across all hosts. Based on this criterion, we identified 434 genes as single host-specific, which clustered into two distinct expression patterns based on their background expression across all hosts: (i) 262 genes were largely un-expressed except within the single host (**Figure 6C; Figure S14B**); and (ii) 172 genes had a low background expression across all hosts with elevated expression in a single host (**Figure 6D; Figure S14C**). Supporting the host-specific *in planta* role for these 434 single host-specific genes, 337 of these genes are expressed only *in planta* with no expression *in vitro*. Of the 97 genes also expressed *in vitro*, 67% had higher expression *in planta* (**Figure S9C**). Together, this indicates that these host-specific genes are mainly functional during colonization on specific hosts. The number of host-specific *B. cinerea* genes was uneven across species, ranging from 83 genes specific to Sunflower to 18 genes specific to Squash (**Figure S14A; Table S10**), suggesting that the degree of host-specific fungal genes varies substantially among hosts.

In addition to these single host-specific genes, we also identified 82 high-entropy genes with elevated expression in two or three hosts (**Figure 6E-F; Figure S15, Table S11**). Among these, 22 showed clustering with host order: Asterales (n = 7), Caryophyllales (n = 9), Fabales (n = 3), and Solanales (n = 3), suggesting limited phylogenetic coherence in host-specific transcriptional responses (**Figure 6E**).

Investigating the role of genetic variation in influencing host-specific genes expression variation showed that they were predominantly shaped by the host. There was some contribution from isolate variation, but solely through host-pathogen interactions (**Figure S8**). This agrees with the expression patterns showing that for each host-specific gene, a single host consistently exhibited higher expression across nearly all 72 *B. cinerea* isolates, relative to other hosts (**Figure 6C–D; Figure S14B–C**). This pattern suggests that these genes are broadly expressed across the diverse isolates in the pathogen population, and they differ because of plastic responses to the host-derived cues, creating the host-specific expression.

To investigate whether host-specific genes organize into co-regulated transcriptional modules, we constructed co-expression networks. Strongly connected networks were observed in Chard, Lettuce, Bean, Cucumber, and Spinach, whereas Cowpea, Squash, Pepper, and Sunflower had smaller networks (**Figure 8**). These 10 networks include a total of 434 genes, of which only 137 had functional annotations. Among the annotated set, several genes were associated with known or putative virulence functions (**Table S10)**. Chard networks included a high number of glycoside hydrolase family genes, along with MFS transporters. In Lettuce, beta-lactamase genes were identified, while the Bean network included *BCIN16G05040* (*BcPKS16*), a polyketide synthase involved in secondary metabolism (Suárez *et al*., 2024). In Spinach, *Bcsdr2*, previously implicated in hyphal growth and pathogenicity on Strawberry and Tobacco, is present (Zhang *et al*., 2024). NADPH oxidase *BcNOxD* was identified in the Sunflower-specific network (Siegmund *et al*., 2015). These findings suggest that *B. cinerea* expresses potential virulence networks in host-dependent manners.

**Figure 8.**
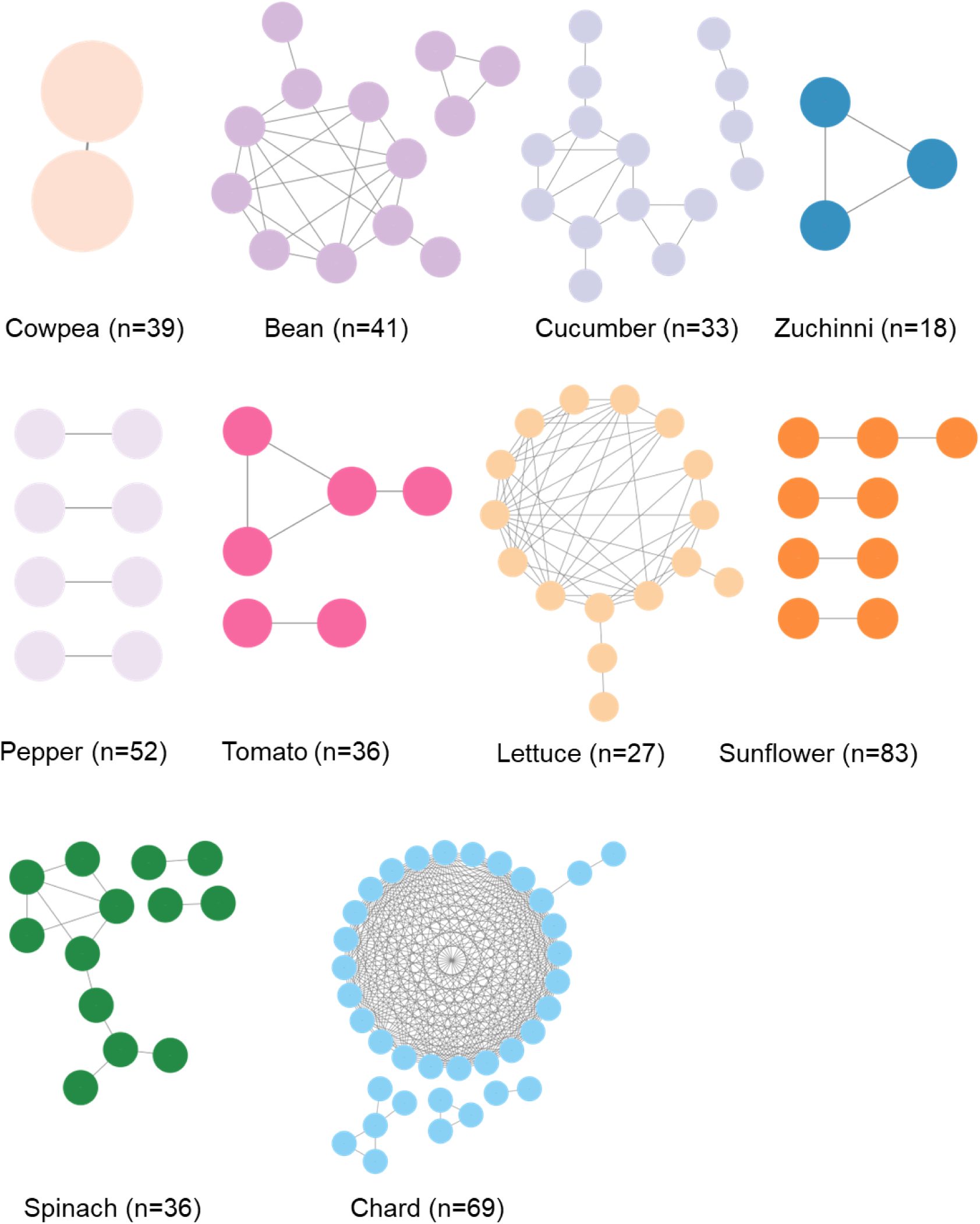
Co-expression networks of *B. cinerea* genes showing host-specific high entropy genes expression in single eudicot hosts. For each host, gene co-expression networks were constructed using Pearson correlation coefficients calculated from gene expression profiles across 72 *B. cinerea* isolates for that host. Nodes represent individual *B. cinerea* genes, and edges denote significant co-expression relationships, defined by the absolute value of the correlation coefficient (|R| ≥ 0.7). The number of genes (n) shown indicates the total number of host-specific genes used as input for that host.

### Similar genomic distribution but differential pressure on general vs host-specific gene sets

To start drawing hypotheses for the different genomic properties and evolutionary pressures influencing the genes associated with host-specific, general lesion, or low entropy expression patterns, we assessed their genomic diversity and chromosomal distribution. First, we analyzed whole-genome sequencing data (Atwell *et al*., 2015) for the isolates and estimated the nucleotide diversity of all genes. This analysis identified that the average non-synonymous (π_N_) and synonymous (π_S_) substitution ratio in protein-coding genes typically occupies a range from 0.15-0.24 with a long right tail (**Figure S16**). This suggests that most genes in the pathogen are under purifying selection. Comparing the general lesion, host-specific high entropy, and constitutive low entropy gene sets to a permutation-based confidence interval revealed that host-specific (high entropy) genes are less constrained by purifying selection than low entropy and general lesion-associated genes (**Figure S16**). This agrees with their host-specific role as any purifying selection would only occur when pathogen is on the specific host for the gene to be expressed and functional. In contrast, the general lesion genes should be under selection when infecting all hosts.

In a number of fungal pathogens, host-specific genes are often clustered within a genome. To test whether the genes associated with host-specific high entropy, general lesion, or low entropy expression patterns exhibited spatial clustering or genomic compartmentalization, we analyzed the chromosomal distribution of these three gene sets. In contrast to other fungal pathogens that display a “two-speed” genome architecture, where adaptive genes are concentrated in gene-sparse, fast-evolving regions (Derbyshire *et al*., 2017; Dong *et al*., 2015; Torres *et al*., 2020), all three gene classes in *B. cinerea* are broadly dispersed across the genome, with only minor evidence of localized clustering (**Figure S17**). This diffuse distribution supports a model in which regulatory plasticity, rather than structural genome compartmentalization or gene presence/absence, underlies host-responsive transcriptional dynamics in *B. cinerea*.

## Discussion

Generalist pathogens such as *B. cinerea* can infect a wide range of plant species, but the molecular strategies enabling this breadth remain elusive. By leveraging a factorial design that incorporates both extensive pathogen diversity and a phylogenetically broad host panel, we identified a modular strategy underpinning generalism that combines general lesion-associated genes linked to primary metabolism and plastic, host-inducible gene modules that respond to specific host environments. This framework supports a model in which specific host-driven transcriptional plasticity in response to the diverse hosts allows *B. cinerea* to infect a broad host range.

### Host specificity via plastic host-inducible genes

Transcriptomic profiling coupled with entropy-based classification identified a subset of genes with strong host-specific expression patterns. These genes are present in nearly all isolates likely explaining why nearly all *B. cinerea* isolates can infect all tested hosts without the requirement of strict host specialization. This model of host specificity distinguishes *B. cinerea* from pathogens such as *F. oxysporum* and *Verticillium dahliae*, where host range is determined by the presence/absence variation of lineage-specific effectors or accessory chromosomes (Chavarro-Carrero *et al*., 2021; Fayyaz *et al*., 2023; Sabahi *et al*., 2021).

While these host-specific genes do not show much presence/absence variation, they do differ across the isolates in their plastic response to the host. This agrees with previous work showing that variation of *in planta* gene expression in *B. cinerea* is to variation in trans regulating networks that influence plasiticty elements (Krishnan *et al*., 2023; Soltis *et al*., 2020). Thus, variation in host specificity appears to derive from genetic polymorphisms in the detection and response to the hosts. evidence points to plant metabolites as potential triggers for these regulatory cascades creating host-specificity. For instance, resveratrol from the grapevine induces *BcLCC2* (Schouten *et al*., 2002), α-tomatine from tomato upregulates glycosyltransferases and RTA1-like transporters (You *et al*., 2024), capsidiol from solanaceous plants activates specific detoxification genes (Kuroyanagi *et al*., 2022), and Brassicaceae glucosinolate derivatives upregulate transporter families (Vela-Corcía *et al*., 2019). This leads to a model where the host specificity of a *B. cinerea* could depend on the phylogenetic distribution of the chemicals ranging from largely family specific like glucosinolates to others that are widely but sporadically present, like saponins or cyanogenic glucosides.

Agreeing with this hypothesis, the above-mentioned detoxification genes fall into the “intermediate entropy” category, neither fully conserved nor strictly host-specific. Their variable expression across hosts would provide a flexible layer, tuned by host cues, allowing *B. cinerea* to adapt infection strategies across rapidly evolving hosts without requiring permanent genetic changes. Such graded responsiveness may be key to enabling generalism while preserving host compatibility. Further work is required to fully understand how *B. cinerea* senses and detects particular hosts.

### Core Metabolism Underlies General Lesion Potential

Alongside host-specific plasticity, we identified 287 genes whose expression was significantly correlated with isolates’ general lesion-forming potential across hosts. These genes formed a co-regulated module enriched in primary metabolic functions and another module with solely members of the *Botcinic acid* biosynthetic genes. Unlike host-inducible genes, these virulence genes were expressed across all hosts but varied in magnitude across isolates. Given that these genes form largely co-regulated modules, it suggests that their variation is also shaped by variation in the trans-regulatory signaling networks controlling these core modules (Krishnan *et al*., 2023). As core metabolism underpins fundamental processes like growth, stress tolerance, resource acquisition and resource allocation, even subtle differences in this module could significantly affect disease severity.

Together, our results position *B. cinerea* as a transcriptionally plastic generalist that combines a flexible, host-inducible repertoire with isolate-specific variation in the regulation of core metabolism. This modular infection strategy enables broad host compatibility while allowing quantitative tuning of virulence. Future work should focus on functional validation of both gene modules and dissection of the regulatory polymorphisms, particularly trans-acting factors, that underlie this plasticity. Expanding this framework to additional hosts, tissues, and environmental conditions, alongside deeper exploration of host-derived cues and host-side dynamics, will enhance our understanding of generalism and its ecological maintenance.

## Materials and Methods

### Plant material and growth conditions

To test how genetic diversity within *B. cinerea* as well as across and within host plant species shapes the disease outcome, host species from a total of eight eudicot orders were included: three Asterids - Apiales, Asterales, and Solanales; one Super-Asterids - Caryophyllales; and four Rosids - Cucurbitales, Fabales, Brassicales, and Malvales (**Figure 1**). Two representative plant species were selected per order, except for the Malvales, which was represented by a single species, *Alcea rosea* (Hollyhock). The species used were: *Petroselinum crispum* (Parsley) and *Apium graveolens* (Celery) from Apiales; *Helianthus annuus* (Sunflower) and *Lactuca sativa* (Lettuce) from Asterales; *Capsicum annuum* (Pepper) and *Solanum lycopersicum* (Tomato) from Solanales; *Spinacia oleracea* (Spinach) and *Beta vulgaris* subsp. *vulgaris* (Chard) from Caryophyllales; *Brassica rapa* (Mustard) and model plant *Arabidopsis thaliana* from Brassicales; *Vigna unguiculata* (Cowpea) and *Phaseolus vulgaris* (Common Bean/Bean) from Fabales; and *Cucumis sativus* (Cucumber) and *Cucurbita pepo* (Squash) from Cucurbitales. These hosts were selected to sample from annual diploid species for which a reference genome is available, and for which there were reports of field infections by *B. cinerea*.

For the leaf infection assays, four genotypes for each plant species were selected to sample intraspecific genetic diversity (**Table S3**). Genotypes were obtained from the USDA-Germplasm Resources Information Network (GRIN), the UC Davis Tomato Genetics Resource Center, the Center for Genetic Resources (CGN) in the Netherlands, and the Michelmore lab from UC Davis. The genotype selection was made in consultation with experts of each crop to maximize genetic diversity based on available information. For clarity, all plant species are referred to as “species” or “hosts” throughout the manuscript.

For the co-transcriptomic study, a subset of five eudicot orders was selected: Asterales and Solanales (Asterids), Caryophyllales (Super-Asterids), and Cucurbitales and Fabales (Rosids), using the same two species per order as described above **(Figure 1)**. To ensure experimental feasibility and enable cross-species comparisons within the broader framework, only one representative genotype per species was used **(Table S3)**. This genotype was chosen based on the lesion data, representing an intermediate lesion development for that host species (**Table S1**).

For both the lesion phenotyping and transcriptomic analysis, all plants were grown in controlled environment chambers in pots containing Sunshine Mix #1 (Sun Gro Horticulture, Agawam, MA, USA). Growth conditions were standardized at 20°C with a 16-hour photoperiod and light intensity of 100–120 μmol m² s¹. Plants were watered every two days with deionized water for the first two weeks, followed by a nutrient solution containing 0.5% N–P–K fertilizer in a 2–1–2 ratio (Grow More 4–18–38).

### *B. cinerea* isolate collection and culture

For all assays, we used 72 *B. cinerea* isolates collected from 14 different host plants and a range of geographical locations (**Table S12**). Our isolate collection contains no significant genetic stratification by either geographic or host species origin (Caseys *et al*., 2021; Soltis *et al*., 2019). All these isolates were previously characterized across eight eudicots (Caseys *et al*., 2021) and on Arabidopsis (Zhang *et al*., 2017; Zhang *et al*., 2019). The 72 *B. cinerea* isolates sample the range of genetic diversity, virulence, and host specificity observed within a larger population of 96 isolates (Atwell *et al*., 2018; Caseys *et al*., 2021).

All the isolates are maintained as conidial suspensions in 60% glycerol at -80 °C for long-term storage. For experiments, spores were grown on potato dextrose agar (PDA) plates by diluting 1:10 (v/v) of glycerol stocks in grape juice and incubated at 21 °C for two weeks.

### *B. cinerea* lesion assays

To enable *B. cinerea* lesion assays comparability across diverse plant species, the standard detached leaf assay was used, using adult leaves as a common organ (Caseys *et al*., 2021; Corwin *et al*., 2016; Denby *et al*., 2004). Detached leaf infections have been shown to correlate well with whole-plant infection assays in various pathosystems (Denby *et al*., 2004; Kirkby *et al*., 2023; Mengiste *et al*., 2003). This approach enabled testing of a large collection of *B. cinerea* isolates while maintaining uniform assay conditions across diverse plant taxa. To ensure consistency in the developmental stage across species, fully expanded mature leaves were sampled during the vegetative phase (4-8 weeks after planting, depending on species-specific growth rates) before the onset of bolting/flowering. This approach minimized ontogenetic variation and ensured comparable leaf maturity across all species used in the experiments. Given the scale of the experiment, it was not possible to introduce developmental or ontogenic variation into the analysis.

Detached leaf assays were performed as previously established (Caseys *et al*., 2021; Denby *et al*., 2004; Soltis *et al*., 2020). In brief, spores were extracted in sterile water, counted using a hemacytometer, and diluted in 50% grape juice to a final concentration of 10 spores/μL. Grape juice, a complex mixture of sugars and micronutrients, was used as the inoculum medium to promote consistent germination across isolates. This alleviates issues where *B. cinerea* has genetic variation impacting germination on single sugar sources (Benito *et al*., 1998; Blakeman, 1975; Clark, 1977; Denby *et al*., 2004). For infection, leaves were placed on trays containing 1 cm of 1% Phytoagar™ (Plant Media), which maintained leaf hydration and physiological activity during the experiment. Leaves were inoculated with 4 μL droplets (40 spores). Grape juice was used as a control. To prevent premature germination during inoculation, spore suspensions were kept on ice and gently agitated to maintain a uniform distribution of spores. Inoculated trays were placed under humidity domes and incubated under constant light. Each isolate × plant genotype combination was replicated three times per experiment in a randomized complete block design. The entire experiment was independently repeated twice, yielding six biological replicates per host-isolate interaction. The lesion area, a quantitative measurement of the host-pathogen interaction, was assessed at 72- and 96-hours post-inoculation (hpi).

For transcriptome analysis, the same setup was used with a single genotype per plant species (**Table S3**). Infection of each isolate on each plant was done as described above (Zhang *et al*., 2017; Zhang *et al*., 2019). Leaf disks surrounding the infection site were harvested at 48 hpi using a cork borer of 1.20 cm diameter, ensuring uniform tissue collection across species. At this timepoint, the lesions were not extensively developed and the majority of the sample was living plant cells. Samples were flash-frozen for RNA-seq library preparation. To confirm infection success, leaves from the same plants, spore batches, and trays were kept at room temperature until 72 hpi to monitor and ensure proper lesion development.

### Lesion measurement and data quality control

Lesion area, a quantitative measure of host-pathogen interaction, was assessed at 72 and 96 hpi. For species that developed large lesions rapidly (Tomato, Sunflower, Lettuce, Arabidopsis, and Mustard), measurements were taken only at 72 hpi to avoid any constraints on lesion development caused by tissue availability. Infection trays were photographed using a Canon T3i camera (18 MP) at a fixed distance and consistent top lighting, which ensured a consistent image quality. Images were analyzed with an R pipeline (Fordyce *et al*., 2018), which converted images to hue/saturation/value (HSV) color space and applied species-specific thresholds to detect lesions. Leaf and lesion masks were generated, manually curated, and the lesion areas were quantified in pixels. A scale included in each image allowed conversion to square millimeters. The dataset comprised 26,718 lesion measurements at 72 hpi and 17,520 at 96 hpi (**Table S1**). Data were filtered to remove technical failures (e.g., droplets without *B. cinerea* growth) as per (Caseys *et al*., 2021).

### Lesion modelin2g

The effect of the host and pathogen genetic variation and experimental factors on lesion area was modeled using linear mixed models implemented in the lme4 package (Bates *et al*., 2015). For each host species, we modeled the lesion area according to the host (genotype), pathogen (isolate), and their interactions while accounting for experimental design.

Linear mixed model: Lesion Area ∼ Genotype + Isolate + Genotype*Isolate + (1|Experiment) + (1|Tray) + (1|Plant)

Here, experiment (experimental replicate), tray (microenvironment containing subsets of leaves within a randomized complete block design), and plant (individual plant identity from which detached leaves were collected) were treated as random effects. Plant genotypes and *B. cinerea* isolates were treated as fixed effects (Corwin *et al*., 2016; Fordyce *et al*., 2018). Given that at the sampling time points, the lesions had not consumed most of the leaf, the leaf area does not influence lesion area development (Caseys *et al*., 2021). Corrected least square means (LS-means) for lesion area for each host genotype x *B. cinerea* isolate combination were extracted from the models and used for downstream analyses (Lenth *et al*., 2019). Variance components were converted to percentages of total variance to facilitate comparisons across models.

To evaluate the relative contributions of host phylogenetic diversity, taxonomic structure, and pathogen virulence to lesion development across host species, we constructed a multi-host meta model using a nested linear model framework. Although the model incorporated plant phylogenetic structure via taxonomic hierarchy, it did not explicitly account for divergence time. Host plants were hierarchically nested by taxonomic levels (Clade > Order > Species > Genotype), and *B. cinerea* isolate was treated as a fixed effect. Interaction terms were included to assess isolate-by-host lineage effects on lesion development.

The multi-host model was specified as: Mean Lesion Area (LS-means) ∼ Isolate + Clade + Clade/Order + Clade/Order/Species + Clade/Order/Species/Genotype + Isolate*Clade + Isolate*Clade/Order + Isolate*Clade/Order/Species.

Hierarchical clustering was performed on standardized LS-means on each host x isolate combination using the complete algorithm, with bootstrap support estimated via the pvclust package (Caseys *et al*., 2021; Kolde, 2019; Suzuki *et al*., 2019). Heatmaps were visualized using the pheatmap package (Kolde, 2019). To minimize bias from missing data, six isolates that failed to sporulate or grow in time for the detached leaf assay of certain species were excluded from this analysis.

### General lesion vs host-dependent lesion

Previous work has shown that it is possible to use lesion area across diverse hosts to estimate the general lesion formation capacity of a *B. cinerea* isolate in addition to measuring the isolate’s lesion formation potential on individual hosts (host-dependent lesion) (Caseys and Kliebenstein, 2025). To estimate the general and host-dependent lesion contributions for the *B. cinerea* isolates, we used linear mixed models. General lesion was estimated by modeling the LS-means of lesion area across all species with the model: Mean Lesion Area (LS-means) ∼ Isolate + (1 | Species) (Caseys and Kliebenstein, 2025). To avoid overfitting and ensure that the general lesion estimate was not biased by performance on the focal species, we implemented a leave-one-species-out approach, whereby the species being evaluated was excluded from the model during its general lesion estimation. This step was critical because including the focal species could inflate the general lesion estimate on that host, especially when regressing host-dependent lesion values against the general lesion, due to circularity. Here, isolate was treated as a fixed effect to estimate isolate-specific lesions across hosts, while fifteen species were treated as a random effect to account for interspecies variability. The average lesion across all hosts for each isolate was obtained as an estimate of general lesion potential.

To estimate each isolate’s lesion-forming potential on individual host species (host-dependent lesion), we fit a separate model for each species: Mean Lesion Area (LS-means) ∼ Isolate + (1 | Genotype). Within each species, the four genotypes were modeled as a random effect to capture within-species genetic variation, while *Botrytis* isolates remained a fixed effect. This model estimated isolate performance on a specific host, adjusting for genotype-level variation.

To quantify deviations in isolate performance from their expected general lesion value, we calculated residuals for each isolate on each host-species from the linear regression: host-dependent lesion ∼ general lesion. These residuals capture the extent to which an isolate’s lesion formation on a given host deviates from its expected general lesion-forming potential. The overall host-specific residual for an isolate was then estimated as the mean of the absolute value of residuals across all species, serving as a quantitative measurement of host specificity. A higher average residual indicated greater variation in isolate performance across hosts, reflecting greater host specificity.

### RNA-Seq library preparation, sequencing, and mapping

*B. cinerea* infected leaf tissues from ten plant species, along with grape juice control, were sampled at 48 hpi for transcriptome analysis as described above. This timepoint was selected based on a pilot RNA-seq experiment conducted at 24, 30, 36, and 48 hpi, which showed that 48 hpi yielded the highest proportion of fungal-mapped reads for all the species. A total of 2190 mRNA libraries were prepared for paired-end sequencing using the Illumina AVITI PE150 platform (DNA Technologies Core, Davis, CA).

RNA-Seq libraries were prepared according to the previous method (Kumar *et al*., 2012) with minor modifications (Zhang *et al*., 2017). Briefly, infected leaves were immediately frozen in liquid nitrogen and stored at -80°C until processing. RNA extraction was conducted by re-freezing samples in liquid nitrogen and homogenizing by rapid agitation in a bead beater, followed by direct mRNA isolation using the Dynabeads mRNA Direct purification kit (Invitrogen). First and second-strand cDNA were produced from the mRNA using an Invitrogen Superscript III kit. The resulting cDNA was fragmented, end-repaired, A-tailed, and barcoded as previously described (Kumar *et al*., 2012). Libraries were size-selected for ∼300 bp and pooled in 96-sample batches for sequencing at the UC Davis Genome Center (DNA Technologies Core, Davis, CA).

Fastq files from individual sequencing lanes were separated by adapter index into individual RNA-seq library samples. Raw RNA-seq reads from individual libraries were subjected to quality control using MultiQC v1.15 to assess overall read quality metrics, including per-base sequence quality and the presence of overrepresented sequences (Ewels *et al*., 2016). Adapter index and low-quality bases were trimmed using Trimmomatic v 0.39 (Bolger *et al*., 2014). Cleaned reads were first mapped to the host reference genome (**Table S13**) using Hisat2 version 2.2.1 with phred33 quality scores and modified alignment parameters to account for mismatches at the read ends (Kim *et al*., 2019). The remaining unmapped reads were subsequently aligned to the *B. cinerea* B05.10 isolate reference genome using the same Hisat2 version and parameters (Van Kan *et al*., 2017). Gene counts were pulled from the resulting SAM files using SAMtools (Li *et al*., 2009) and converted to BAM files. Custom R scripts were used to summarize counts across gene models to reduce the overrepresentation of genes with multiple splice variants.

### *B. cinerea* gene expression analysis

Raw read counts for *B. cinerea* genes were processed and normalized using R (v4.3.1) and the edgeR package (v3.42.4) (Robinson *et al*., 2009). To remove lowly expressed genes while preserving biologically relevant, isolate- or host-specific transcripts, we applied a minimal expression filter, retaining genes with a count-per-million (CPM) value of > 1 in at least two samples per host using the cpm function in edgeR. In this case, CPM refers solely to the reads that map to *B. cinerea* in the sample. This threshold minimizes background noise without excluding lineage-specific or conditionally expressed genes. Gene counts passing this filter were normalized using the Trimmed Mean of M-values (TMM) method. Normalized CPM values were extracted with library size correction and subsequently averaged across biological replicates for each isolate–host combination, resulting in one expression value per gene per group.

To estimate *B. cinerea* transcript abundance in infected tissue, we calculated the percentage of total reads (host plus pathogen) that mapped to *B. cinerea* genes (% *B. cinerea* reads) for each RNA-Seq sample. This was done by dividing the number of reads mapped to *B. cinerea* genes by the total mapped reads per sample and multiplying by 100:

% *B. cinerea* reads = (*B. cinerea* mapped reads / Total mapped reads) × 100.

This value was computed independently for each plant species and tested as a proxy for fungal biomass.

To test how the *B. cinerea* transcripts vary across the diverse host plants, the isolate variation, and their interaction, we fitted a linear mixed model for each transcript.

Linear mixed model: CPM ∼ (1 | *B. cinerea* isolate) + (1 | Host) + (1 | Isolate:Host)

where CPM denotes the normalized counts per million for each transcript, isolate represents 72 *B. cinerea* isolates, host represents the 10 eudicot species, and the interaction term accounts for host– pathogen interactions. Models were run separately for the full dataset as well as subsets at the clade and order levels to explore broader phylogenetic patterns. Modeling was not performed at the individual species level due to the presence of only one genotype per species.

For each transcript, broad-sense heritability (*H^2^*) was calculated as the proportion of total variance explained by genetic sources (isolate, host, and their interaction) (Krishnan *et al*., 2023). The resulting *H²* values were visualized using ternary density plots generated with the ggtern package (Hamilton and Ferry, 2018) in R (v4.3.1), which illustrate the relative contribution of each source to transcriptomic variation.

### Shannon Entropy Calculation and Classification of Genes

Shannon entropy is commonly used to distinguish constitutively expressed genes from condition- or tissue-specific genes (Ameri and Lewis, 2021; Heintzman *et al*., 2009; Zhang *et al*., 2006). In the context of plant–pathogen interactions, we applied Shannon entropy to quantify variability in *B. cinerea* gene expression across host species using the BioQC package (Ameri and Lewis, 2021). Genes with low entropy show similar expression across the hosts and were classified as conserved. In contrast, high entropy genes are highly variable across hosts and were used to classify host-specific genes, where gene expression was limited to one or a few specific hosts.

To focus on the expression patterns observed across hosts and reduce noise due to the genetic diversity across the 72 *B. cinerea* isolates, we calculated the mean of CPM-normalized expression values within each host species. To validate this method, entropy was also calculated for all 72 isolates, and the resulting gene ranks were concordant across methods. The genes were categorized into low, intermediate, and high entropy using Jenks’ natural breaks optimization function (getJenksBreaks in BAMMtools) (Jenks, 1967). Based on Jenks clustering, 589 genes had high entropy (H> 3.25), 10,828 genes were intermediate (3.25>H<0.56), and one gene had low entropy (H< 0.0030).

To account for the range and distribution of entropy values and allow identification of the host(s) associated with high-entropy genes, gene expression values were log-transformed [log₂(CPM + 1)] and z-score normalized across host species for each gene individually. A gene was considered highly expressed in a particular host if its z-score exceeded +1, indicating expression at least one standard deviation (SD) above its mean.

Genes with z-scores > +1 SD in only one host were classified as single-host-specific (n = 434), while those with z-scores > +1 SD in two to three hosts were labeled as multi-host-specific (n = 82). Seventy-three genes initially assigned to the high entropy group by Jenks clustering but exhibiting low inter-host variance were excluded based on the z-score > +1 criterion. Each single-host-specific gene was assigned to the host with the highest z-score.

### General lesion-associated genes

To identify *B. cinerea* genes associated with the general lesion estimate across diverse host species, we fitted the linear model for each gene:

General lesion ∼ Gene expression + Host + Gene expression x Host.

This model tests the relationship between fungal gene expression (CPM value) and general lesion, while accounting for host identity and gene-by-host interaction effects. *P*-values associated with each model term were corrected for multiple testing using the Benjamini–Hochberg false discovery rate (FDR). Genes with FDR-adjusted *p-*values below 0.05 for the main gene expression term in the model (i.e., independent of host or interaction effects) were considered as general lesion-associated genes and used for further analysis.

### Co-Expression Network Analysis of High-Entropy, Low-Entropy, and General Lesion-Associated Genes

Gene co-expression networks were constructed using Pearson correlation coefficients to evaluate potential coordination among different gene classes: host-specific high-entropy genes, conserved low-entropy genes, and general lesion-associated genes.

For the high-entropy networks, only genes with elevated expression (z-score > +1) in a particular host were included in that host’s network. For example, a gene showing the highest expression (z-score > +1) in Chard was included exclusively in the Chard-specific network, and its expression across the 72 *B. cinerea* isolates infecting Chard was used to compute pairwise correlations. This approach ensured that each host-specific network captured gene co-regulation patterns uniquely associated with transcriptional responses induced by that particular host.

In contrast, the low-entropy and general lesion-associated gene networks were constructed using expression data across all 720 isolate–host combinations. Given that only a single low-entropy gene was detected by Jenks’ clustering, we included 500 genes with the lowest entropy values to low-entropy network. The general lesion-associated network included 287 genes.

For all networks, Pearson correlation coefficients were computed for all gene pairs, and edges were retained if the absolute correlation was ≥ 0.7, a threshold chosen to reflect strong co-expression while minimizing noise (Mukaka, 2012). Networks were built using the igraph package in R and exported as edge lists and node degree tables for visualization in Cytoscape v3.10. This resulted in a total of 10 host-specific co-expression networks, in addition to the low-entropy and a general lesion-associated gene networks.

### Comparison of *In Planta* and *In Vitro B. cinerea* Transcriptomes

To estimate *in vitro* gene expression and compare to *in planta* estimates, we analyzed publicly available RNA-seq datasets of the reference isolate *B05.10* grown in potato dextrose broth (PDB), a media with the same sugar composition as the PDA used in our infection assays. Only control or mock samples were used from three projects: PRJNA1056687 (6–48 h; (Lu *et al*., 2025)), PRJNA955032 (0 h; (You *et al*., 2024)), and PRJNA1173356 (0–24 h), collectively capturing gene expression over multiple growth timepoints. The reads were aligned and normalized with the same pipeline and parameters as mentioned above for *in planta* analysis.

To compare *in vitro* and *in planta* expression, we used only *B05.10* isolate data across ten host species (*in planta*). Comparisons were performed separately for three gene sets: (1) 500 conserved low-entropy genes, (2) all 434 single-host-specific high-entropy genes, and (3) 287 general lesion-associated genes. For each gene within each set, average normalized expression (log₂ [CPM + 1]) was calculated separately across all *in vitro* and *in planta* samples. The absolute difference between average *in planta* and *in vitro* expression values was used to classify each gene into one of three expression categories: (1) Similar expression in both *in vitro* and *in planta* (absolute difference ≤ log_2_ unit); (2) Higher expression *in planta* (absolute difference >1 log_2_ unit *in planta*); and (3) Higher *in vitro* (absolute difference >1 log_2_ unit *in vitro*).

### Inference of the strength of purifying selection

To estimate the selective pressure on *B.cinerea* genes, we analyzed the whole-genome sequencing data (PRJNA525902) available for the isolates (Atwell *et al*., 2015). In short, the Illumina reads were the B05.10 reference genome assembly ASM83284v1 (Van Kan et al., 2017) with bwa mem (Li, 2013). Single nucleotide polymorphisms (SNPs) were extracted with Freebayes (Garrison and Marth, 2012). The rate of non-synonymous (π_N_) and synonymous (π_S_) substitutions in protein-coding genes were estimated in SNPGenie (Nelson *et al*., 2015). To provide a confidence interval of π_N_/π_S_ values to compare the three gene sets, we also analyzed 100 permutations of 500 random genes.

## Data, Materials, and Software Availability

All data supporting the findings of this study are included in the main article and/or the Supplementary Information. The RNA-seq data have been deposited in the NCBI Sequence Read Archive (SRA) under BioProject ID **[PRJNA1217477]**, and will be made publicly available upon publication.

## Acknowledgements

This work was supported by the NSF award IOS 2020754 to DJK. We thank all members of the Kliebenstein lab for their invaluable assistance and help throughout the course of this large-scale experiment.

## Authors contributions

D.J.K.: designed research; R.S., A.J.M., C.T., J.M.M., K.S., L.F., and C.C.: performed research; R.S.: analyzed data; and R.S.: Writing—original draft, D.J.K and C.C.: Writing—review and editing.

## Supplementary Figures

**Figure S1.**
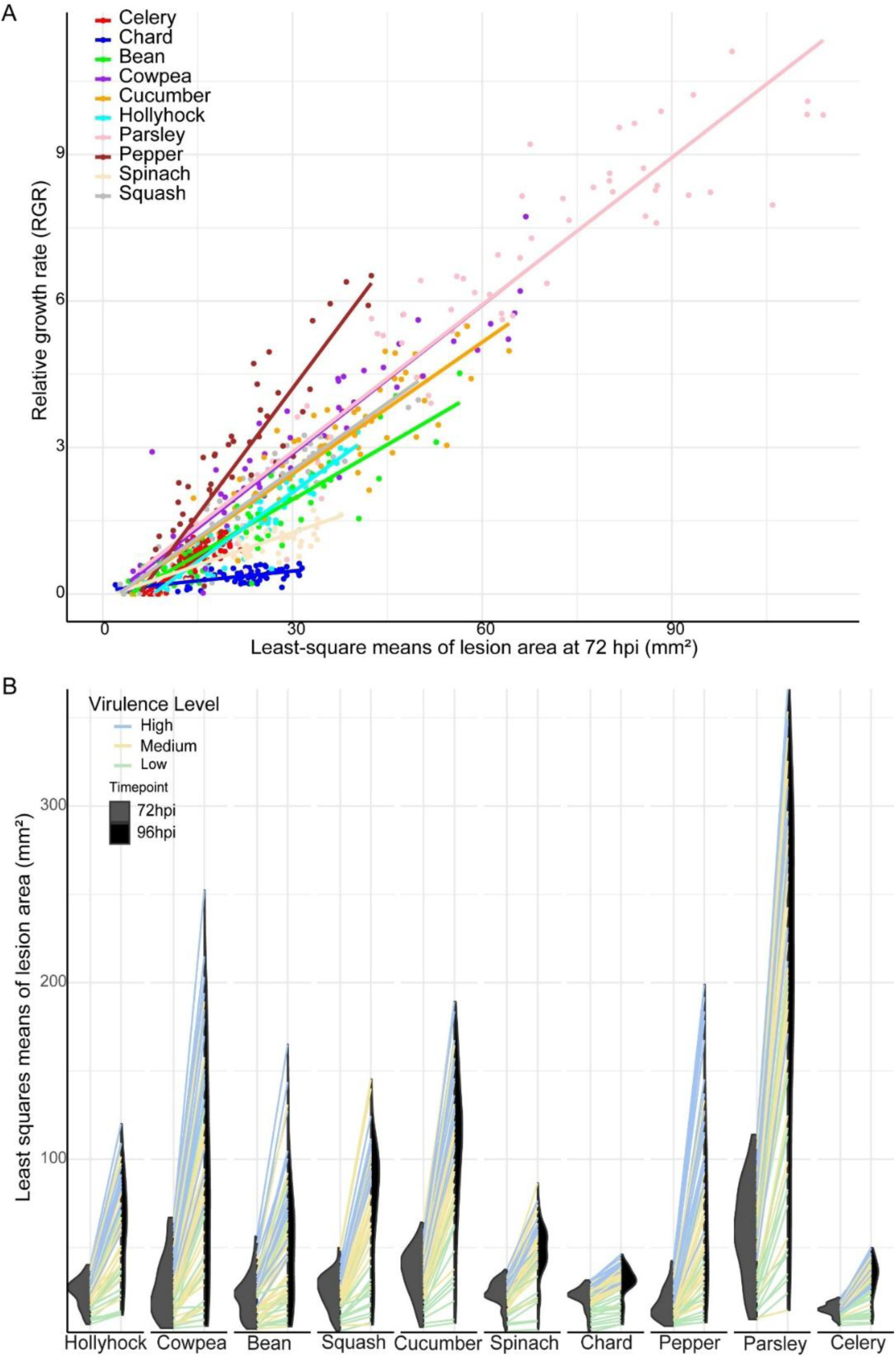
Lesion progression dynamics of *B. cinerea* from 72 to 96 hours post-inoculation (hpi) across 10 eudicot host species. **(A)** Scatterplot showing the linear relationship between mean lesion area at 72 hpi (LS-mean) and the relative growth rate (RGR) of lesions from 72 to 96 hpi across 10 host species. Each point represents a *B. cinerea* isolate, colored by host species. Solid lines indicate linear regressions fitted separately for each host. The subset of 10 species used for co-transcriptomes was selected for this time-course analysis. **(B)** Half violin plots showing lesion progression from 72 (gray) to 96 (black) hpi across 10 eudicot host species. Lines connect data points for each *Botrytis cinerea* isolate, which are classified into three virulence categories based on their mean lesion area across all hosts at 72 hpi: high (blue), medium (yellow), and low (green) virulence. Only 10 host species are shown for this time-course analysis due to constraints on lesion development caused by tissue availability.

**Figure S2.**
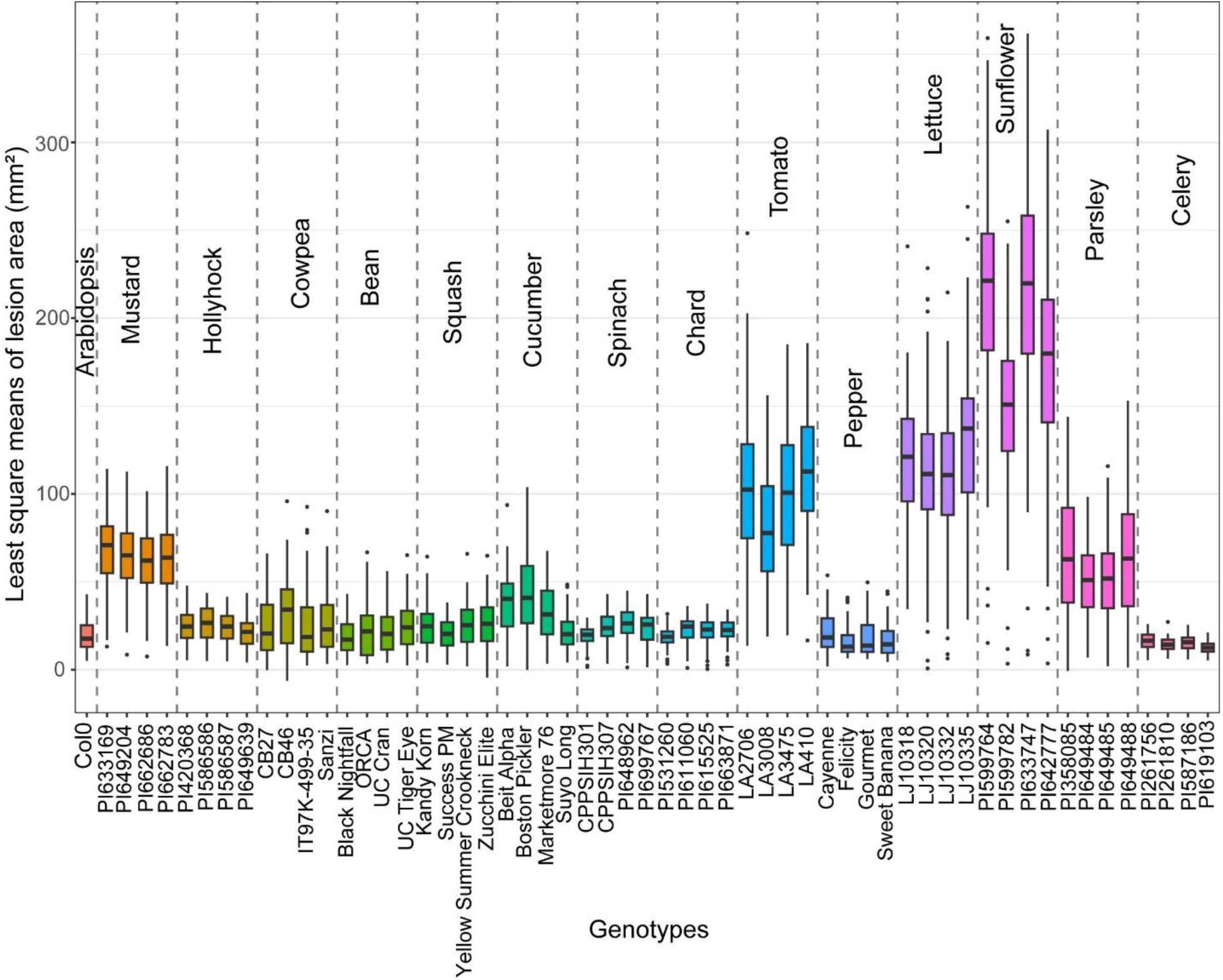
Lesion development and host susceptibility across eudicot species. **(B)** Boxplots showing least squares mean (LS-mean) lesion area (mm²) at 72 hpi for individual genotypes across 15 eudicot plant species. Genotypes are grouped by species and separated by dotted lines. The species name is labeled once per group above the corresponding set of boxplots, and genotype names are shown along the x-axis. Species are arranged according to their phylogenetic order and clade classification. Arabidopsis is represented by a single genotype (Col-0), while all other species are represented by four genotypes each.

**Figure S3.**
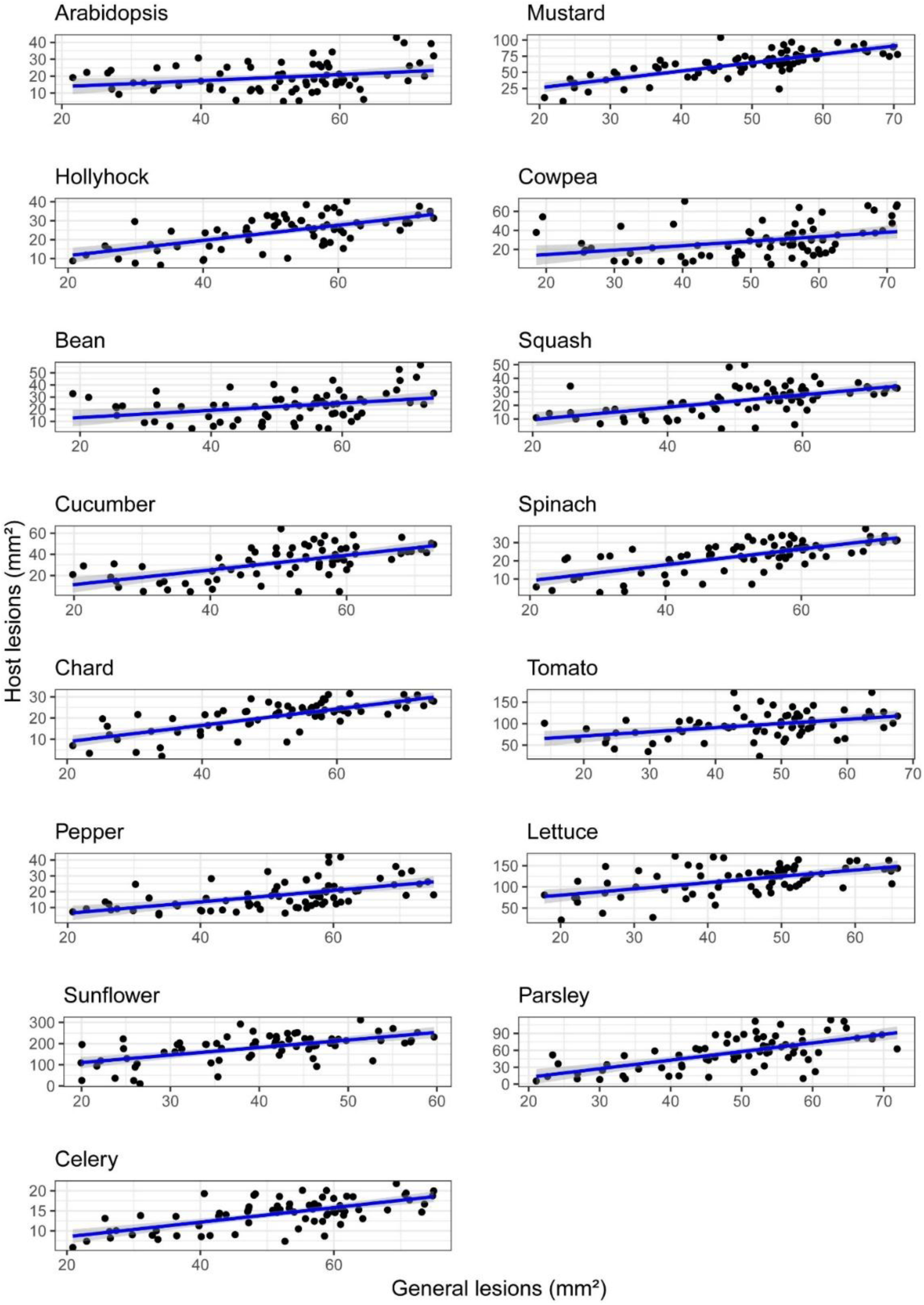
Relationship between general lesion and host lesion size across 15 eudicot hosts. Scatterplots illustrate the relationship between general lesion size (x-axis; average lesion size across all hosts, excluding the focal host) and species-specific lesion size (y-axis; lesion size measured on the focal host) for each of the 15 eudicot plant species. Each point represents a single *B. cinerea* isolate. Blue lines indicate linear regression fits for each host. Corresponding *R^2^* and *p* values are provided in **Table S2**.

**Figure S4.**
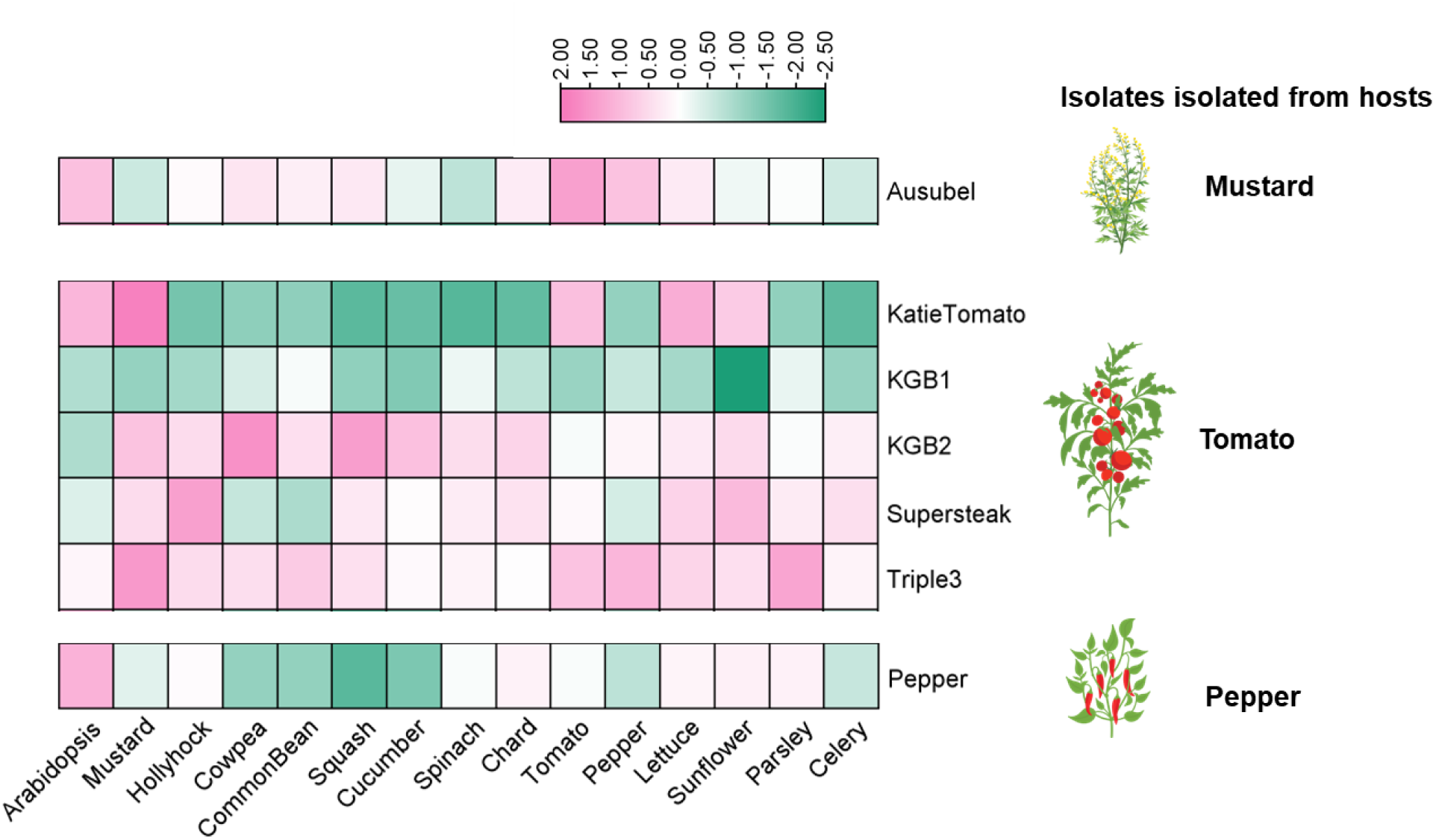
Source-host specialization patterns of *B. cinerea* isolates. Z-scaled heatmap showing lesion size across 15 eudicot plant hosts for seven *B. cinerea* isolates, each originating from a specific host. Isolates include five tomato-derived isolates (*KatieTomato*, *KGB1*, *KGB2*, *Supersteak*, and *Triple3*), one mustard-derived isolate (*Ausubel*), and one pepper-derived isolate (*Pepper*). Rows represent individual *B. cinerea* isolates, and columns represent host plant species. Color values represent z-scaled responses across hosts, with pink indicating higher-than-average values and green indicating lower-than-average values.

**Figure S5.**
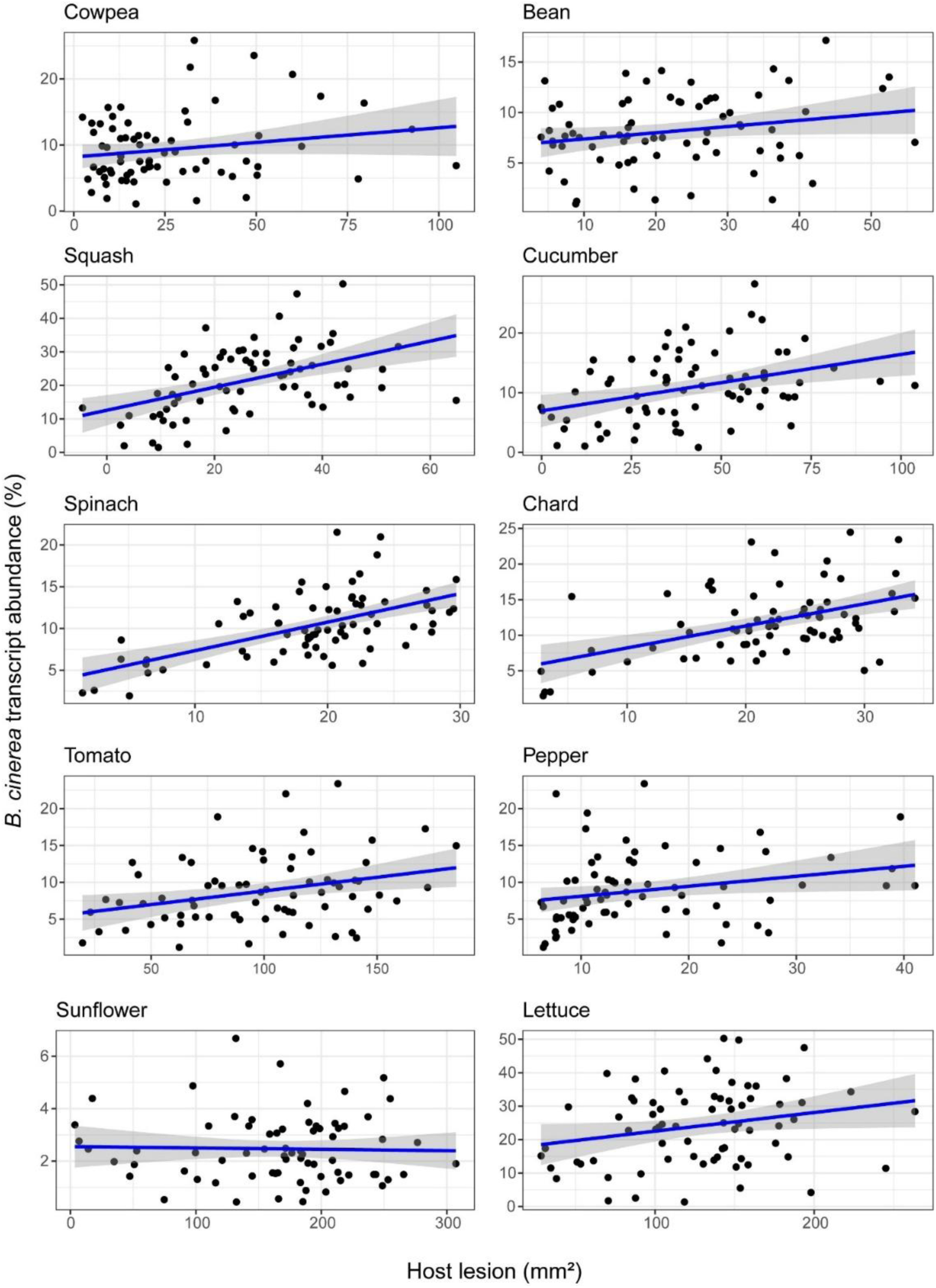
Relationship between *B. cinerea* transcript abundance and host lesion across 10 eudicot hosts. Scatterplots show the relationship between lesion size at 72 hpi on each host species (host lesion; x-axis) and early *B. cinerea* transcript abundance at 48 hpi (y-axis). Each point represents a single *B. cinerea* isolate. Blue lines indicate linear regression fits for each host. Corresponding R² and *p*-values are provided in **Table S4**.

**Figure S6.**
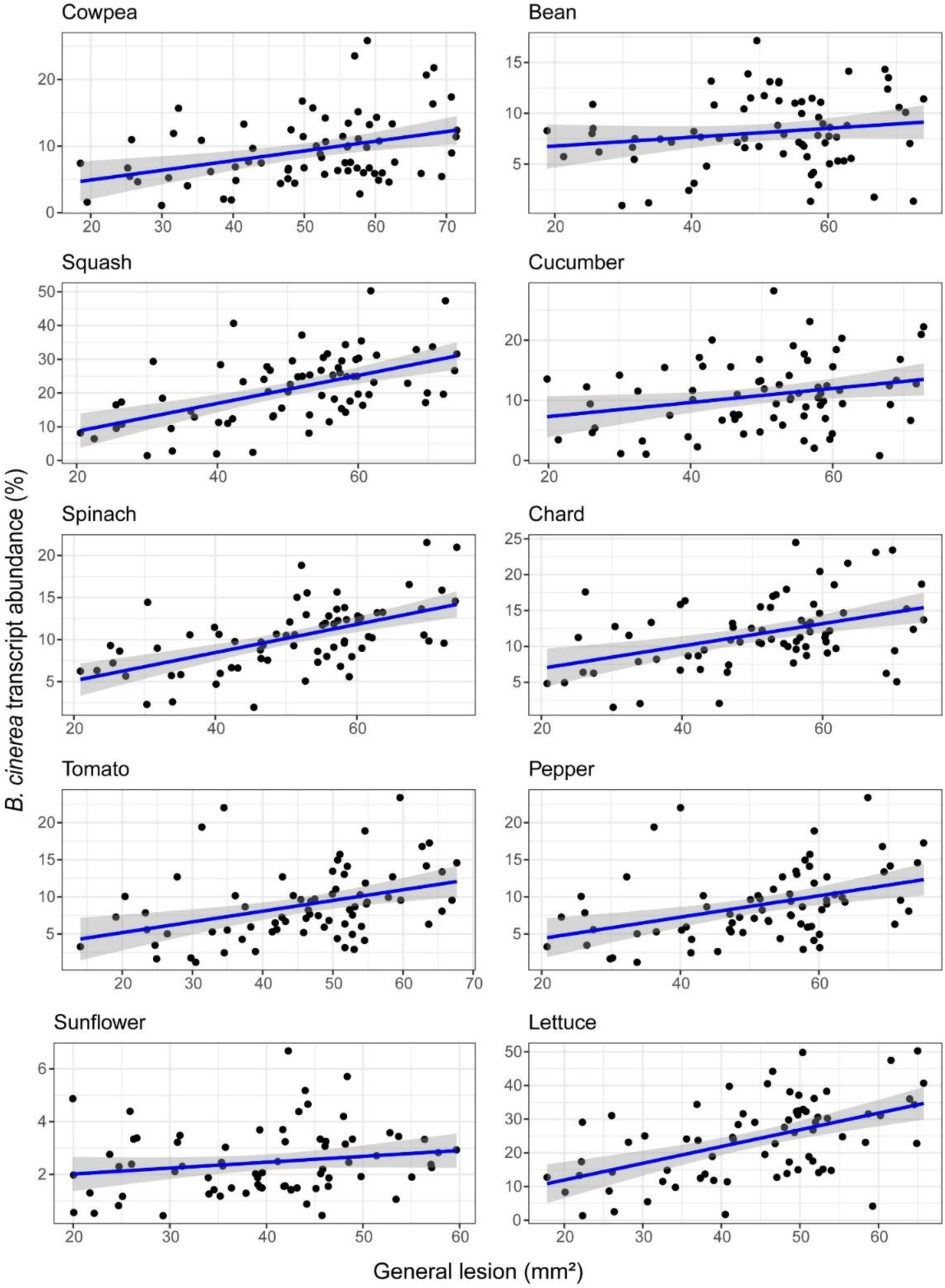
Relationship between *B. cinerea* transcript abundance and general lesion across 10 eudicot hosts. Scatterplots show the relationship between general lesion at 72 hpi (mean lesion area across all host species; x-axis) and early *B. cinerea* transcript abundance at 48 hpi (y-axis) for each host species. Each point represents a single *B. cinerea* isolate. Blue lines indicate linear regression fits. Corresponding R² and *p*-values are provided in **Table S4**.

**Figure S7.**
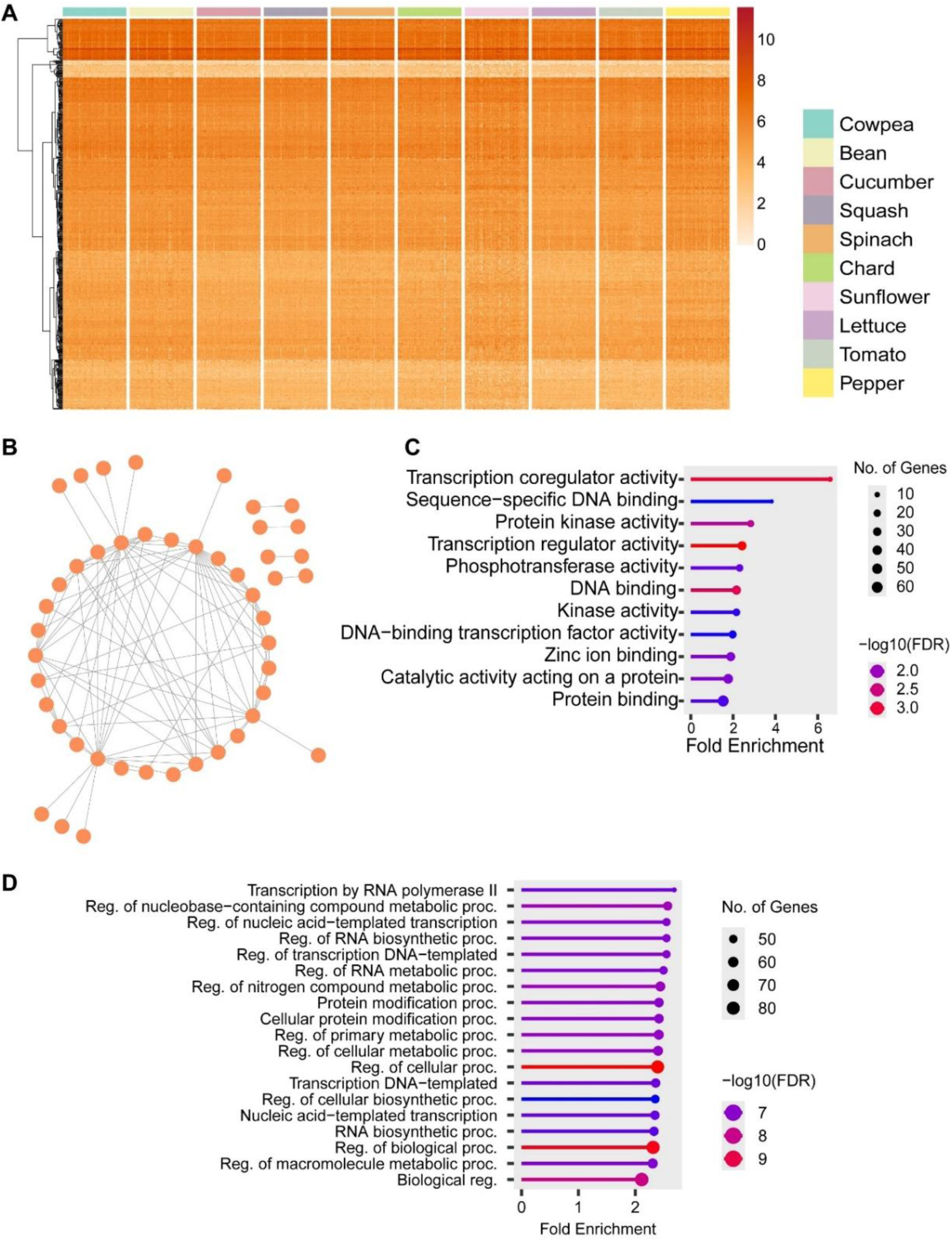
Expression patterns, co-expression network, and functional enrichment of 500 low-entropy *B. cinerea* genes. **(A)** Heatmap showing log₂ (CPM + 1) expression levels of low-entropy genes across 72 *B. cinerea* isolates infecting 10 eudicot plant hosts. Columns represent host species (color-coded), and rows represent conserved *B. cinerea* genes hierarchically clustered by expression profile. **(B)** Co-expression network of the top 500 conserved genes, constructed from pairwise Pearson correlation analysis using the absolute correlation coefficient (|r| ≥ 0.7) across the entire *B. cinerea* transcriptome dataset (72 isolates × 10 hosts = 720 samples). Nodes represent individual genes; edges represent significant co-expression relationships. **(C–D)** Gene Ontology (GO) enrichment analysis for the 500 low-entropy genes. Dot plots show enriched **(C)** molecular function and **(D)** biological process categories, with dot size representing the number of genes per term and color indicating significance level (–log₁₀[FDR]), generated using ShinyGO.

**Figure S8.**
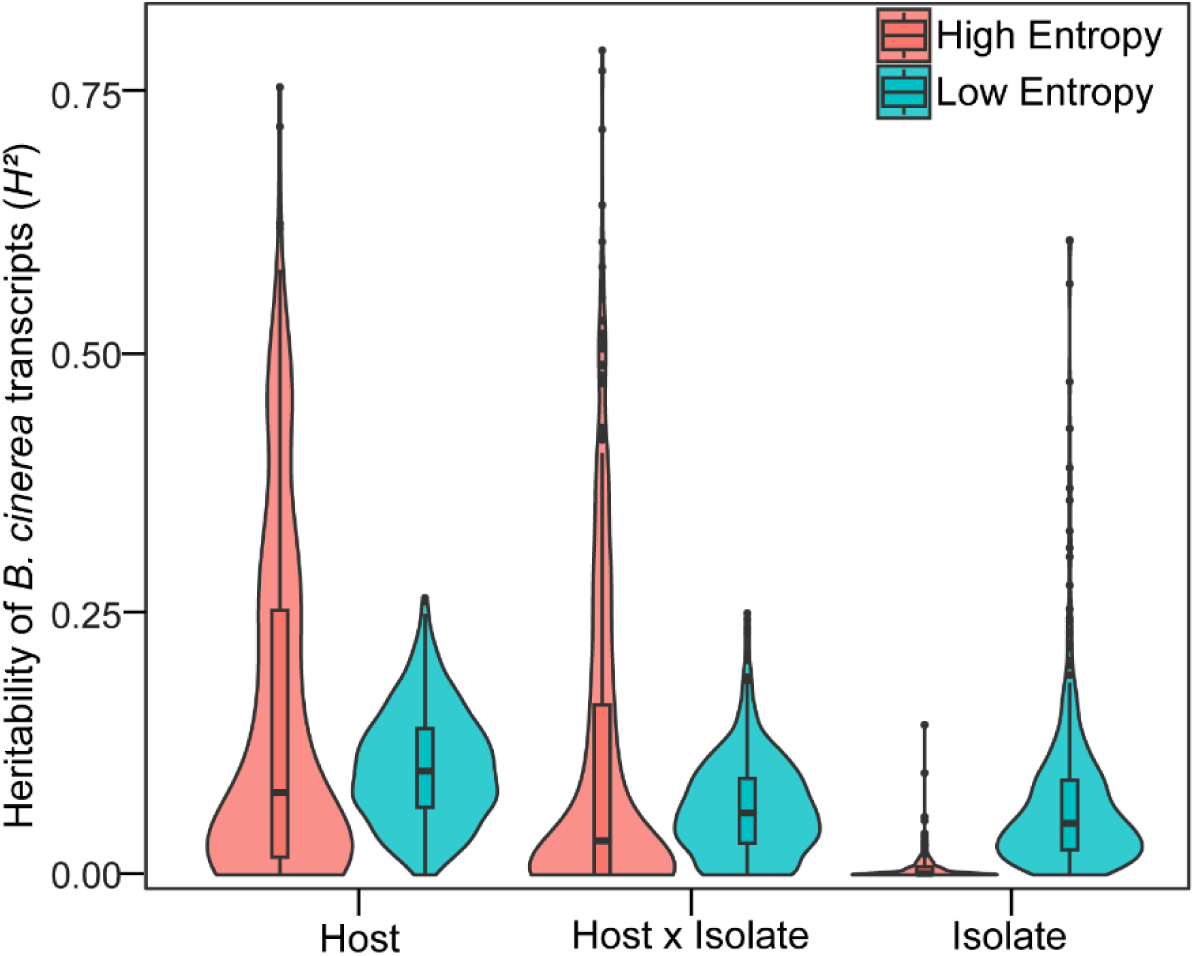
Distribution of broad-sense heritability (*H²*) of *B. cinerea* transcripts explained by isolate, host species, and their interaction for high- and low-entropy genes. Violin and box plots show the distribution of *H²* for *B. cinerea* gene expression across 72 isolates infecting 10 eudicot host species. Heritability estimates were partitioned into three components based on linear mixed-effects models: isolate: genetic variation attributable to 72 *B. cinerea* isolates; host: variation attributable to 10 eudicot host plant species, and their respective interactions.

**Figure S9.**
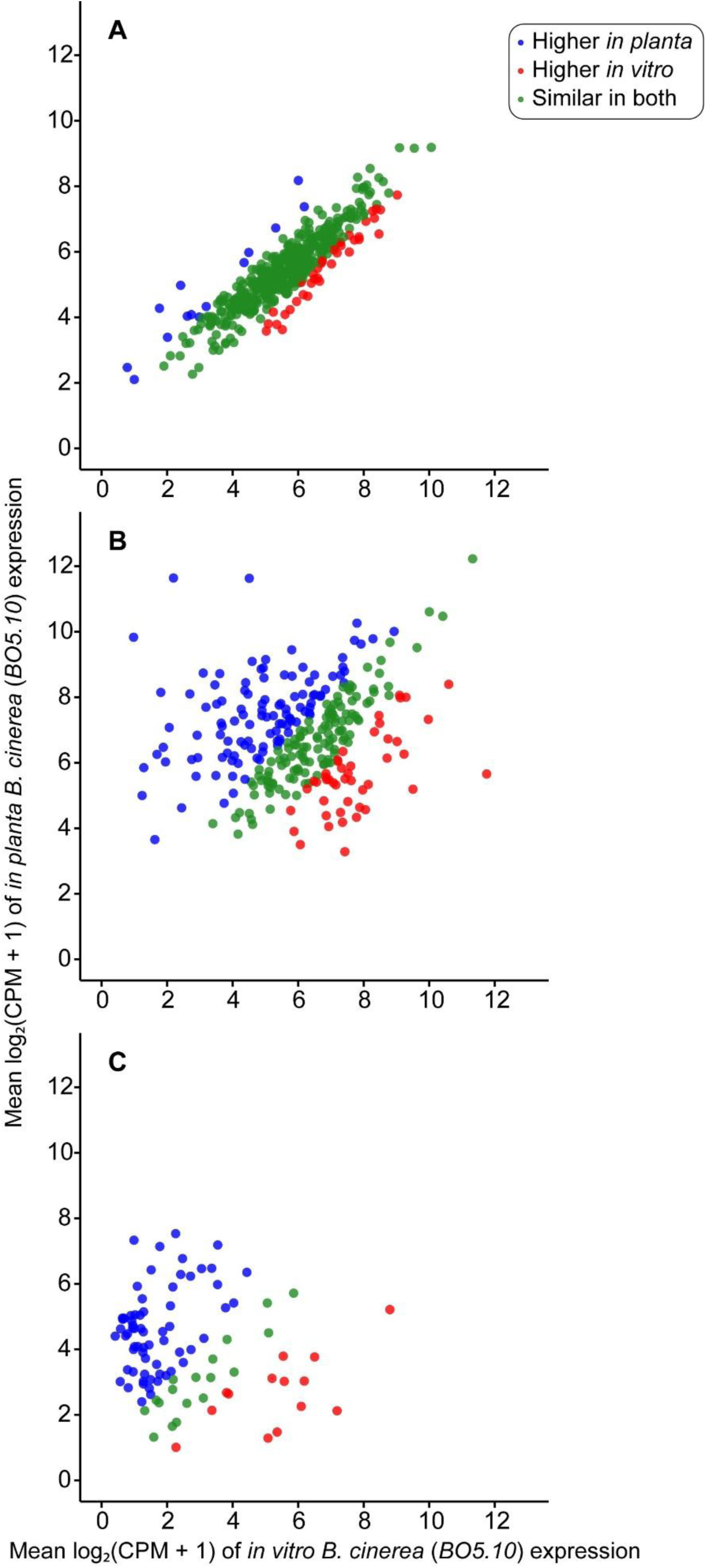
Comparison of *B. cinerea* (*BO5.10* isolate) transcript accumulation *in planta* and *in vitro*. Scatterplots show average expression (log₂ [CPM + 1]) of *B. cinerea* genes measured *in planta* (across 10 eudicot hosts) versus *in vitro* (PDB media) for the *B05.10* isolate. Each point represents a single gene, colored by relative expression bias: **(A)** 500 low-entropy (conserved) genes. **(B)** 287 general lesion–associated genes. **(C)** 434 high-entropy (host-specific) genes; only 97 showed detectable expression *in vitro*.

**Figure S10.**
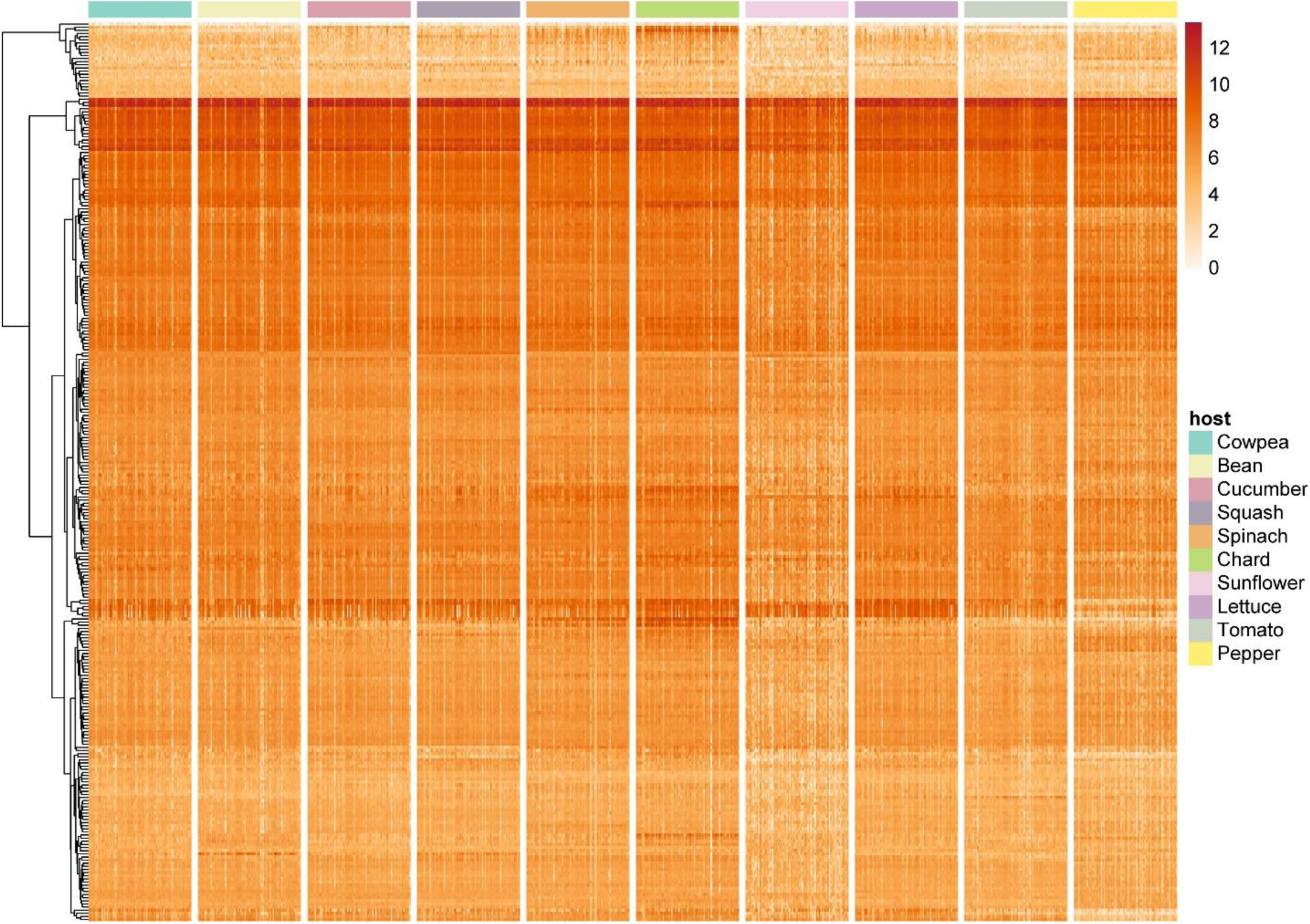
Expression patterns of general lesion–associated *B. cinerea* transcripts across 10 eudicot species. Heatmap of log₂ (CPM + 1) expression values for the same 287 genes across all 72 *B. cinerea* isolates infecting 10 eudicot species. Columns represent host species (color-coded), and rows represent individual *B. cinerea* genes clustered by expression profile. The color scale bar reflects log₂ (CPM + 1) values.

**Figure S11.**
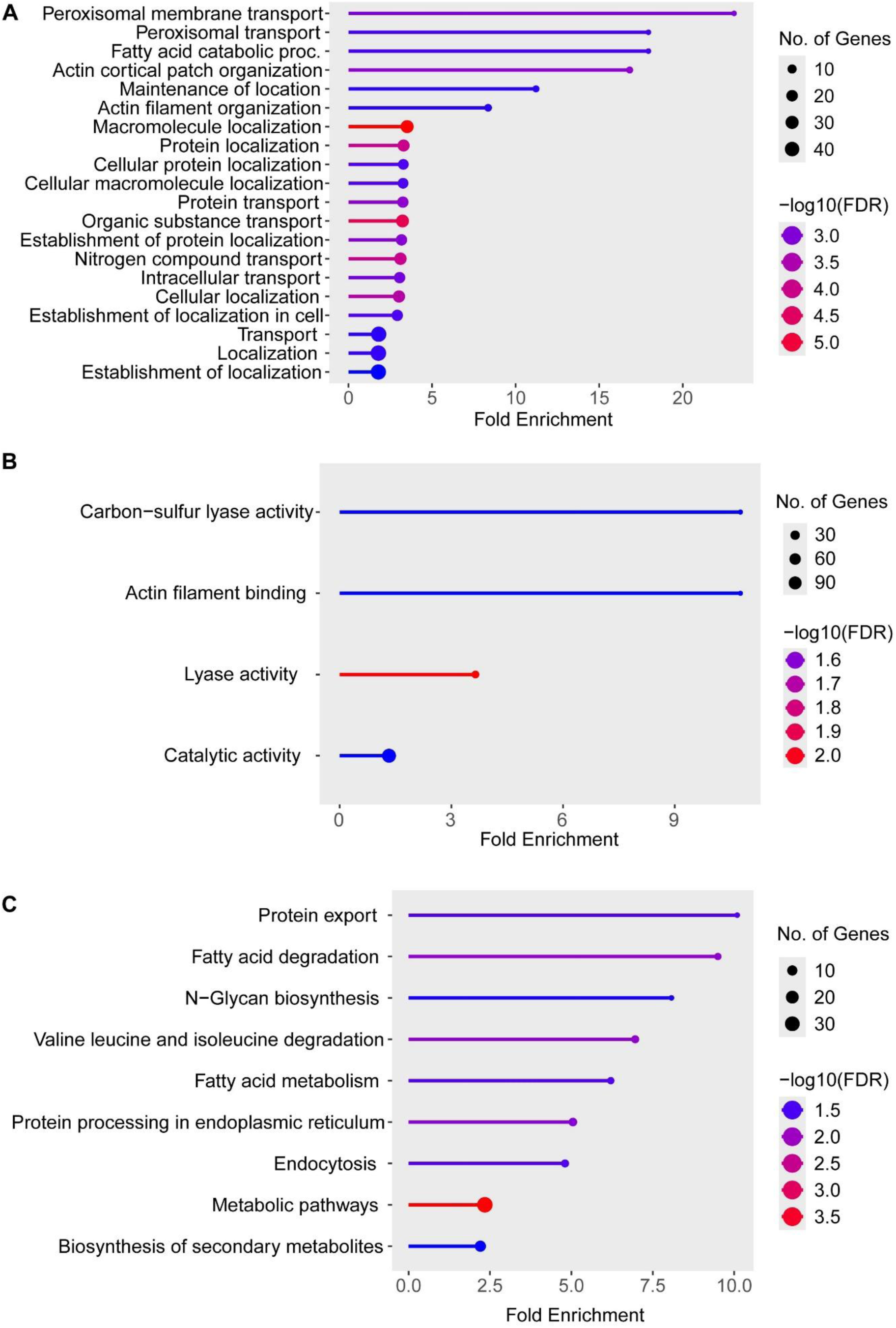
Functional enrichment analysis of general lesion–associated *B. cinerea* genes. Gene Ontology (GO) and KEGG enrichment analyses were performed for 287 *B. cinerea* transcripts significantly associated with general lesion size using Shiny GO. Significantly enriched categories are shown for **(A)** GO Biological Processes (BP), **(B)** GO Molecular Functions (MF), and **(C)** KEGG pathways. Dot plots display enriched terms, with dot size corresponding to the number of genes annotated in each category and dot color representing statistical significance (– log₁₀(FDR)).

**Figure S12.**
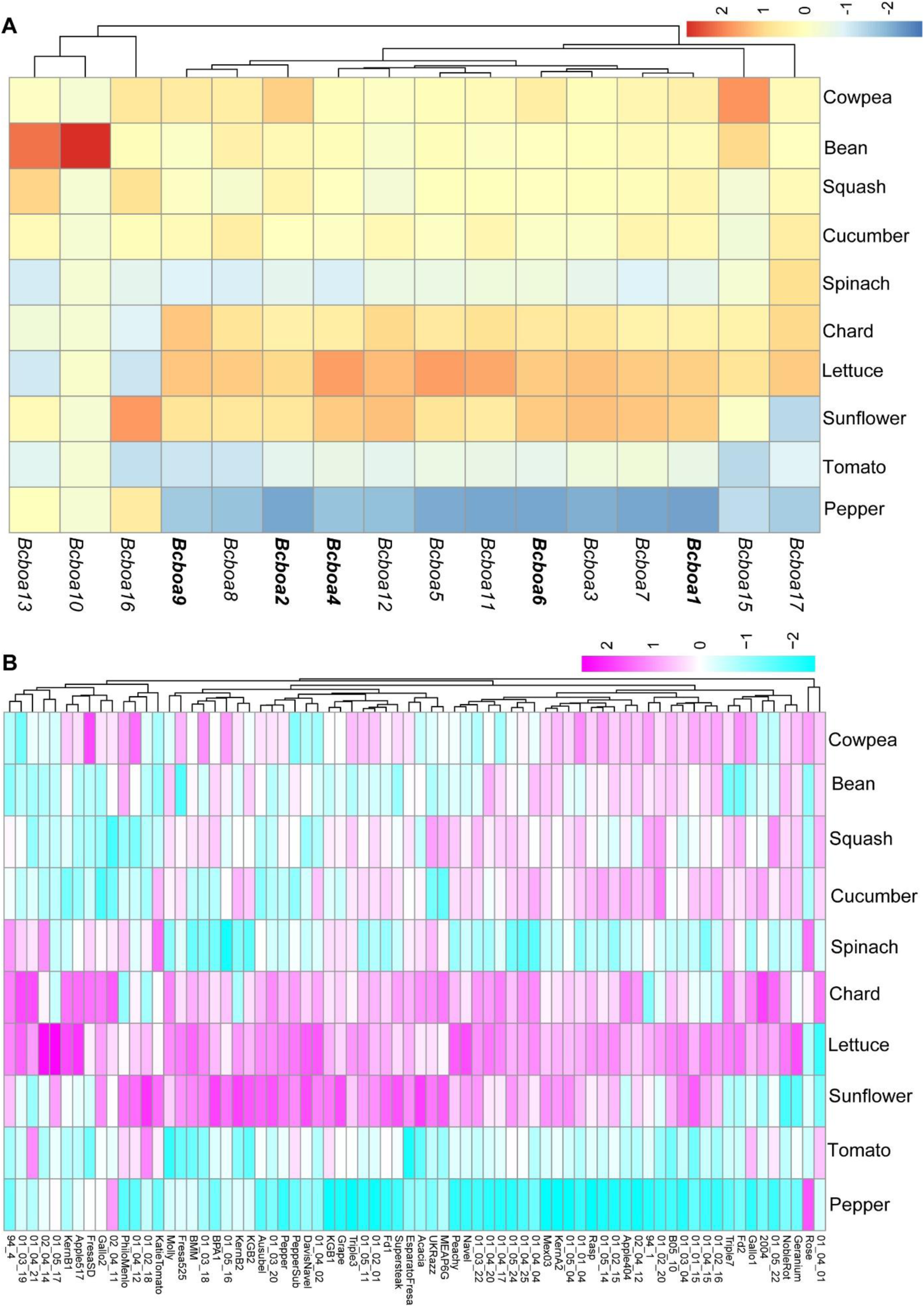
Expression patterns of *B. cinerea* botcinic acid (BOA) cluster genes across 10 eudicot species. **(A)** Heatmap showing z-scaled expression of individual BOA cluster genes (columns; *Bcin01g00010*-*Bcin01g00160*) averaged across all isolates infecting each host species (rows). Genes included are based on the full annotated BOA biosynthetic cluster. Hierarchical clustering was applied to both genes and host species. Genes highlighted in bold (*Bcboa1*, *Bcboa2*, *Bcboa4*, *Bcboa6*, *Bcboa9*) are members of a co-expressed submodule identified in general lesion-associated network analysis. **(B)** Heatmap showing z-scaled expression of the BOA cluster averaged at the gene-set level for each isolate–host combination. Columns represent individual *B. cinerea* isolates, grouped by host species (rows).

**Figure S13.**
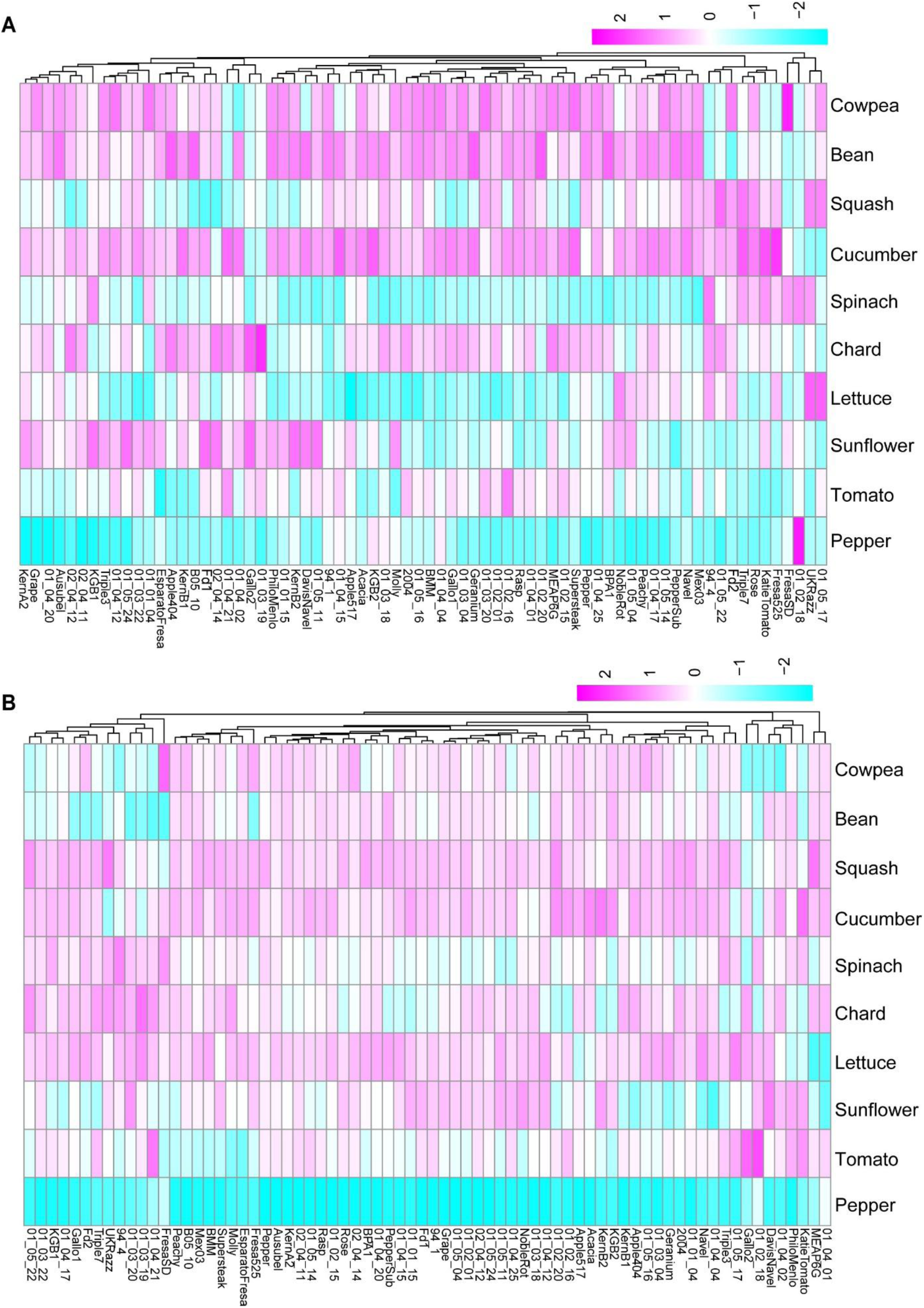
Expression patterns of canonical virulence genes across 72 *B. cinerea* isolates infecting 10 eudicot host species. (A) Z-scaled heatmap showing expression of all genes in the *Botrydial* (*BOT*; *Bcin12g06370*–*Bcin12g06410*) biosynthetic cluster, averaged at the gene-set level across isolate–host combinations. (B) Z-scaled heatmap showing expression of the polygalacturonase gene *Bcpg1* (*Bcin14g00850*) across the same isolate–host matrix. Columns represent individual isolates; rows are grouped by host species. Hierarchical clustering was applied to isolate axis.

**Figure S14.**
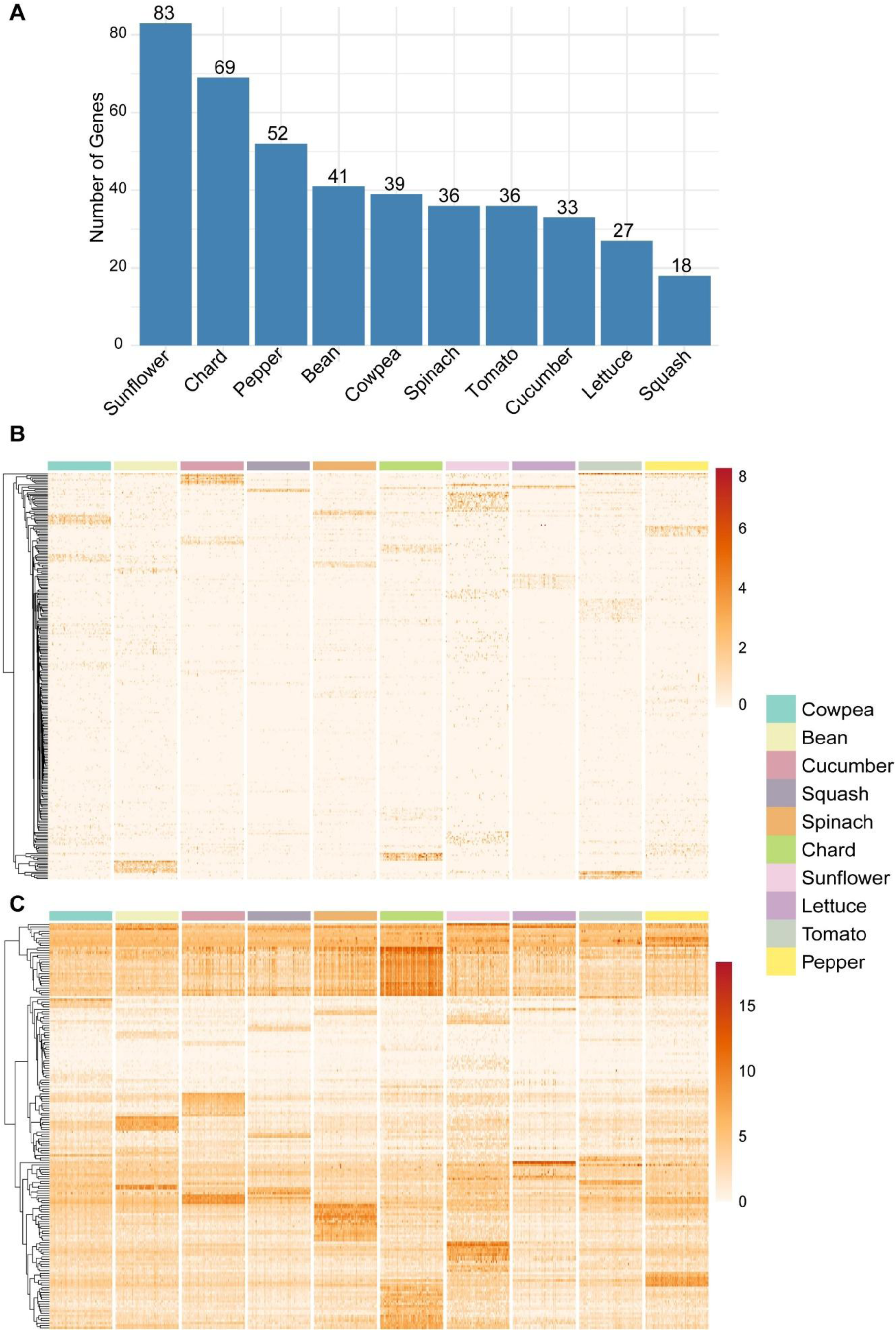
Distribution and expression patterns of single host–specific genes across *B. cinerea* isolates infecting 10 eudicot host species. **(A)** A gene was defined as single host-specific if its expression in one host was ≥ 1 standard deviation (SD) above its mean expression across all hosts. Each gene was assigned to the host in which it showed ≥1 SD higher expression. The bar plot shows the number of single host-specific genes identified per host. **(B-C)** Heatmap of log₂ (CPM + 1) expression values for single-host-specific genes, shown across all 72 *B. cinerea* isolates with (**B)** 262 genes expressed in only a few hosts while **(C)** 172 genes expressed across hosts, but with expression ≥ 1 SD in one host compared to others. Columns represent host species (color-coded), and rows represent individual genes hierarchically clustered by expression profile.

**Figure S15.**
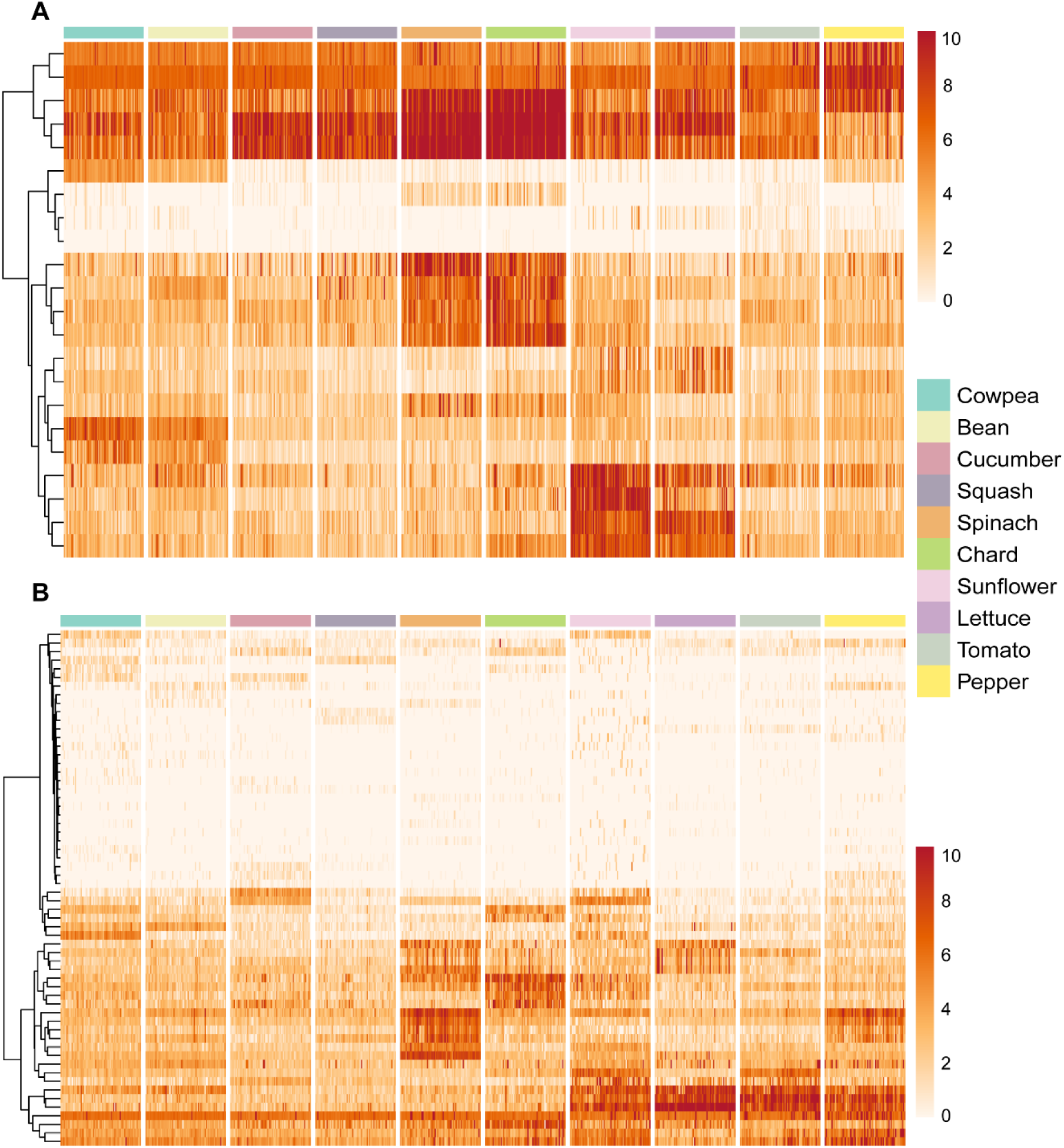
Multi-host-specific gene expression profiles across *B. cinerea* isolates infecting 10 eudicot hosts. **(A-B)** Heatmaps of log₂ (CPM + 1) expression values for genes showing moderate host specificity, defined as having expression ≥1 standard deviation above the gene’s mean in only two or three host species. All 72 *B. cinerea* isolates are shown across all 10 host. **(A) G**enes (n=22) with high expression in phylogenetically related hosts, suggesting order-specific regulatory responses. **(C)** Genes (n=60) with high expression in two or three phylogenetically unrelated hosts. Columns represent host species (color-coded), and rows represent individual *B. cinerea* genes hierarchically clustered by expression profile.

**Figure S16:**
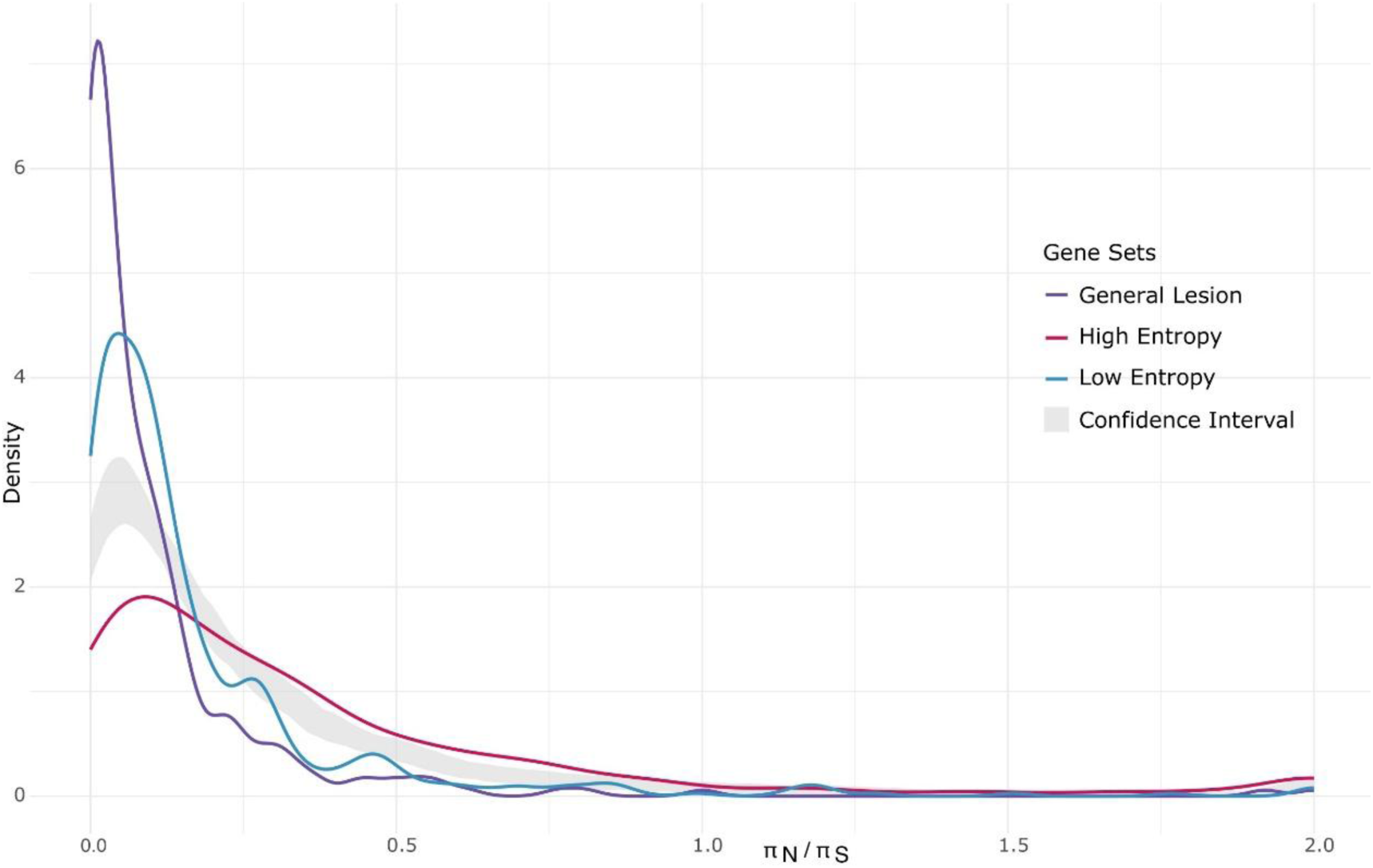
Density distribution of non-synonymous to synonymous nucleotide diversity ratios (πN/πS) for three *Botrytis cinerea* gene sets: general lesion-associated, host-specific high-entropy, and constitutive low-entropy genes. The shaded grey area represents the 95% confidence interval of 100 permutations comprising 500 random genes.

**Figure S17.**
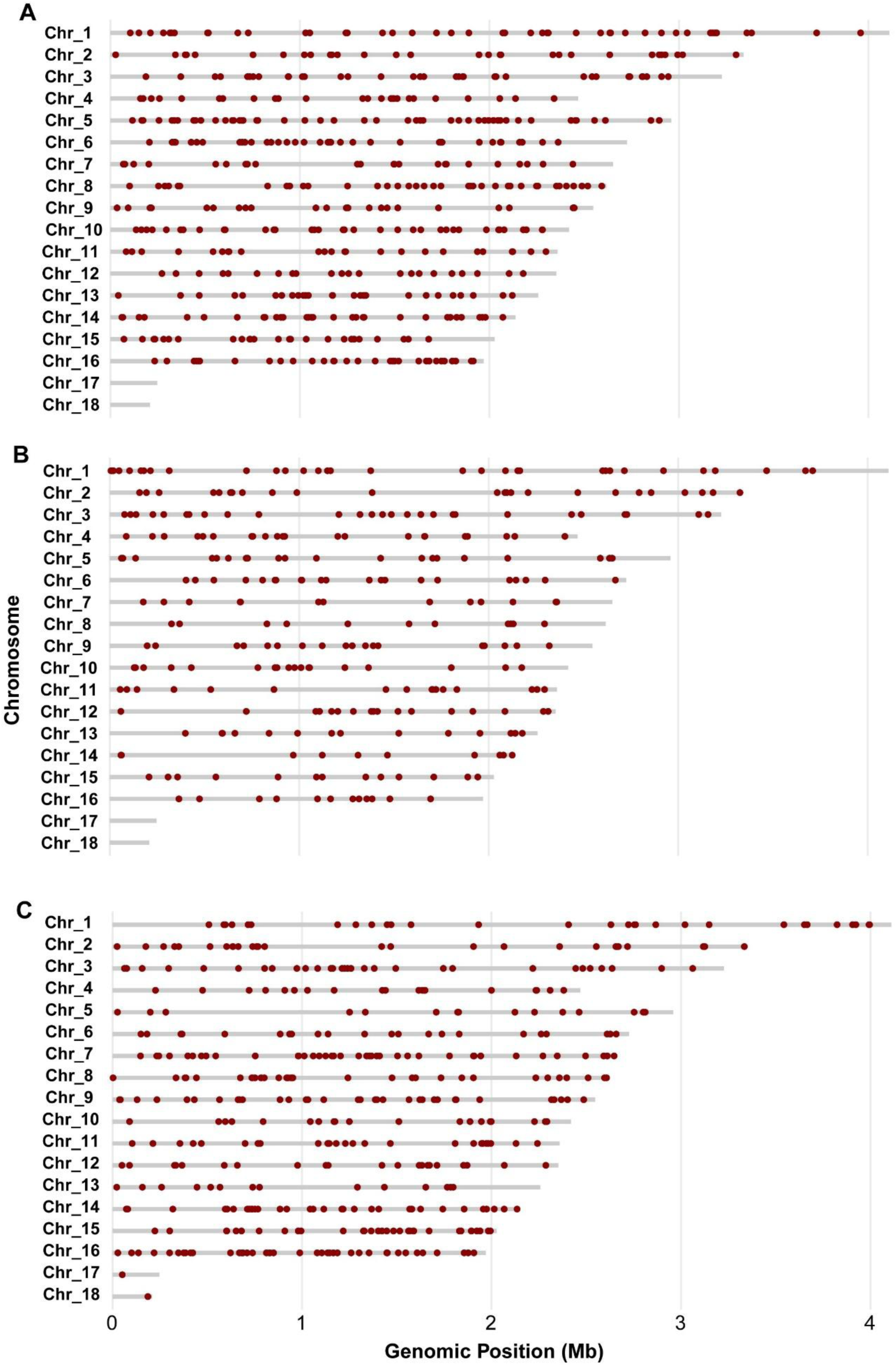
Genomic distribution of entropy-classified and general lesion–associated *B. cinerea* genes across 18 chromosomes. **(A)** Low-entropy genes. **(B)** General lesion-associated. **(C)** High-entropy genes. Plots were generated using ShinyGO. The x-axis indicates chromosomal position in megabase pairs (Mbp); each red dot represents an individual gene.

