## Supplementary Tables for "Combined generalist and host-specific transcriptional strategies enable host generalism in the fungal pathogen *Botrytis cinerea*": Table_S2_GLvsHL_r2.docx

**Table S2: Coefficient of determination (R²) and p-values from linear regression models testing the relationship between host-dependent lesion and general lesion across *B. cinerea* isolates for each species.**

| **Species** | **R^2^** | **P value** |
| --- | --- | --- |
| Arabidopsis | 0.066 | 0.0175 |
| Mustard | 0.586 | 2.97e-15 |
| Hollyhock | 0.391 | 4.27e-09 |
| Cowpea | 0.111 | 0.00247 |
| Bean | 0.101 | 0.00377 |
| Squash | 0.321 | 1.29e-07 |
| Cucumber | 0.35 | 2.62e-08 |
| Spinach | 0.458 | 4.02e-11 |
| Chard | 0.538 | 1.45e-13 |
| Tomato | 0.157 | 0.000374 |
| Pepper | 0.278 | 1.16e-06 |
| Lettuce | 0.276 | 1.29e-06 |
| Sunflower | 0.329 | 1.03e-07 |
| Parsley | 0.405 | 1.11-09 |
| Celery | 0.422 | 3.93e-10 |
