## Supplementary Tables for "Combined generalist and host-specific transcriptional strategies enable host generalism in the fungal pathogen *Botrytis cinerea*": Table_S4_BT_.docx

**Table S4:** Coefficient of determination (*R²*) and *p*-values from linear regression models testing the relationship between *B. cinerea* transcript abundance (BT) and lesion size across isolates for each host species. The models include: (1) Host-dependent lesion (HL) ~ BT, and (2) General lesion (GL) ~ BT.

|  | **HL vs BT** | | **GL vs BT** | |
| --- | --- | --- | --- | --- |
| **Species** | **R^2^** | **P value** | **R^2^** | **P value** |
| Cowpea | 0.022 | 0.114 | 0.127 | 0.00122 |
| Bean | 0.035 | 0.0632 | 0.013 | 0.167 |
| Squash | 0.211 | 2.96e-05 | 0.283 | 8.9e-07 |
| Cucumber | 0.121 | 0.00159 | 0.055 | 0.0261 |
| Spinach | 0.333 | 6.72e-08 | 0.314 | 1.86e-07 |
| Chard | 0.225 | 1.52e-05 | 0.169 | 0.000195 |
| Tomato | 0.082 | 0.00886 | 0.132 | 0.000993 |
| Pepper | 0.046 | 0.0396 | 0.148 | 0.000491 |
| Lettuce | 0.041 | 0.0482 | 0.255 | 3.59e-06 |
| Sunflower | -0.014 | 0.818 | 0.019 | 0.255 |
