## Supplementary Tables for "Combined generalist and host-specific transcriptional strategies enable host generalism in the fungal pathogen *Botrytis cinerea*": TableS8_HLGL_lowentropy.docx

**Table S8:** Spearman correlation coefficients (ρ) and *p*-values for low-entropy, general lesion-associated, and high-entropy genes with host-dependent lesion for each eudicot species.

|  | **Low-entropy genes** | | **General-lesion associated genes** | | **Host-specific high-entropy genes** | |
| --- | --- | --- | --- | --- | --- | --- |
| **Species** | **Spearman rho** | **Spearman *p*** | **Spearman rho** | **Spearman *p*** | **Spearman rho** | **Spearman *p*** |
| Chard | 0.327 | 0.00525 | 0.691 | 0 | 0.544 | 1.24e-06 |
| Bean | -0.195 | 0.101 | 0.129 | 0.281 | 0.252 | 0.0329 |
| Cowpea | -0.142 | 0.234 | 0.225 | 0.0578 | -0.083 | 0.485 |
| Cucumber | 0.098 | 0.41 | 0.58 | 1.72e-07 | -0.356 | 0.0023 |
| Lettuce | -0.057 | 0.633 | 0.243 | 0.0403 | 0.064 | 0.594 |
| Pepper | -0.006 | 0.96 | 0.15 | 0.208 | 0.011 | 0.929 |
| Spinach | 0.028 | 0.818 | 0.583 | 1.40e-07 | -0.157 | 0.187 |
| Squash | 0.072 | 0.546 | 0.476 | 2.95e-05 | 0.038 | 0.749 |
| Sunflower | -0.013 | 0.914 | 0.17 | 0.156 | 0.194 | 0.105 |
| Tomato | 0 | 1 | 0.211 | 0.0778 | -0.091 | 0.452 |
